# Temporal regulation of prenatal embryonic development by paternal imprinted loci

**DOI:** 10.1101/700948

**Authors:** Qing Li, Yuanyuan Li, Qi Yin, Shuo Huang, Kai Wang, Liangchai Zhuo, Wei Li, Boran Chang, Jinsong Li

## Abstract

*H19* and *Gtl2* are paternal imprinted genes that are pivotal for prenatal embryonic development. Meanwhile, mouse nongrowing oocytes and sperm- or oocyte-originated haploid embryonic stem cells (haESCs) carrying both *H19* and *IG*-DMR (differentially DNA-methylated region) deletions (DKO) that partially mimic paternal imprinting of *H19*-*Igf2* and *Dlk1*-*Dio3* can be employed as sperm replacement to efficiently support full-term embryonic development. However, how *H19*-DMR and *IG*-DMR act together to regulate embryonic development is still largely unknown. Here, using androgenetic haESC (AG-haESC)-mediated semi-cloned (SC) technology, we showed that paternal *H19*-DMR and *IG*-DMR are not essential for pre-implantation development of SC embryos generated through injection of AG-haESCs into oocytes. *H19*-DMR plays critical roles before 12.5 days of gestation while *IG*-DMR is essential for late-gestation of SC embryos. Interestingly, we found that combined deletions of *H19* and *H19*-DMR can further improve the efficiency of normal development of SC embryos at mid-gestation compared to DKO SC embryos. Transcriptome and histology analyses revealed that *H19* and *H19*-DMR combined deletions rescue the placental defects. Furthermore, we showed that *H19*, *H19*-DMR and *IG*-DMR deletions (TKO) give rise to better prenatal and postnatal embryonic development of SC embryos compared to DKO. Together, our results indicate the temporal regulation of paternal imprinted loci during embryonic development.

## INTRODUCTION

In the 1970s, many groups attempted to induce mouse parthenogenetic (PG) development, however, the PG embryos could not survive after 10 days of gestation (Graham, 1970; Kaufman, 1973; Tarkowski, 1975). In 1984, two elegant embryological experiments by construction of mouse embryos carrying only paternal or maternal genome provided direct evidence to demonstrate nonequivalence of parental genome, the concept of genomic imprinting (McGrath and Solter, 1984; Surani et al., 1984). Genomic imprinting, an epigenetic mechanism that results in parent-of-origin specific gene expression, plays essential roles in the regulation of mammalian embryonic and extraembryonic development (Barlow and Bartolomei, 2014; Ferguson-Smith, 2011).

Among the 151 reported murine imprinted genes (http://www.mousebook.org/imprinting-gene-list), the *H19*-*Igf2* locus, the first identified imprinted region, is critical for normal prenatal growth and placentation and regulated by differentially DNA methylation (*H19*-DMR) (Arney, 2003; Bartolomei et al., 1991; Cleaton et al., 2014; DeChiara et al., 1991). *H19*-DMR, located 2 to 4 kb upstream from the start of transcription start site (Thorvaldsen et al., 1998; Tremblay et al., 1997), is essential for establishing the pattern of imprinting by which *H19*, a non-coding RNA, is exclusively expressed from the maternal chromosome (Bartolomei et al., 1991) and *Igf2* is only active paternally (DeChiara et al., 1991). On the maternal chromosome, the DMR is hypomethylated and bound by the zinc-finger protein CCCTC-binding factor (CTCF) (Bell et al., 1999), leading to the establishment of an insulator act as a barrier to influence the neighboring *cis*-acting elements and silence *Igf2* on the maternal allele (Bell and Felsenfeld, 2000; Hark et al., 2000). On the paternal chromosome, the DMR is hypermethylated and not bound by CTCF, thus allowing downstream enhancers to activate paternal *Igf2* gene (Bell and Felsenfeld, 2000; Hark et al., 2000). *H19* knock-out mice are viable, fertile, but present an overgrowth phenotype (Leighton et al., 1995). In contrast, overexpression of *H19* was deleterious to embryo development after embryonic day 14 (E14) (Brunkow and Tilghman, 1991). Deletion of *Igf2* leads to fetal and placental growth restriction (Constância et al., 2002; DeChiara et al., 1990), whereas overexpression of the *Igf2* by imprint relaxation leads to placental and fetal overgrowth (Eggenschwiler et al., 1997). Interestingly, paternal deletion of *H19*-DMR, although with *H19* activation and decreased *Igf2* expression, results in overall normal growth in the mouse (Leighton et al., 1995; Thorvaldsen et al., 1998). Taken together, the *H19*-*Igf2* locus influences embryonic development probably through regulating the expression of *Igf2*, which plays important roles in embryonic and placental development (Burns and Hassan, 2001; Morrione et al., 1997; Sibley et al., 2004).

The *Dlk1*-*Dio3* is another essential imprinted region for embryonic development, in which, *Gtl2* and *Dlk1* are reciprocally expressed (*Gtl2* is expressed from maternal allele, while *Dlk1* is expressed exclusively from the paternal allele) (Schmidt et al., 2000; Takada et al., 2000) and controlled by intergenic germline-derived (*IG*)-DMR (Lin et al., 2003; Takada et al., 2002). Deletion of *IG*-DMR from maternal chromosome leads to loss of imprinting (LOI) in this locus and causes embryonic lethal after E16, but paternal deletion does not change the imprinting state of the region, leading to normal embryonic development in the resultant mouse (Lin et al., 2003).

With the identification of the imprinted genes, scientific efforts have been made to remove the paternal barriers that prevent the normal development of mouse bi-maternal embryos.

Interestingly, PG embryos, which contained one set of chromosome from a neonate-derived non-growing (ng) oocyte with imprinting free and the other from a fully grown (fg) oocyte, successfully developed to normal-sized fetus with a well-developed placenta on day 13.5 of gestation, 3 days longer than previously reported for parthenogenetic development (Kono et al., 1996). Furthermore, using the same system, when ng oocytes harboring a 3-kilobase (kb) deletion covering *H19* transcription unit, the reconstructed parthenotes could develop to day 17.5 of gestation (Kono et al., 2002). Moreover, when ng oocytes carrying a 13-kb deletion including both *H19* and *H19*-DMR, a small ratio of the parthenotes could develop to term and grow to adulthood (Kono et al., 2004). Surprisingly, the birth rate of the reconstructed bi-maternal embryos was greatly improved when both *H19*-DMR and *IG*-DMR were deleted in the ng oocytes, indicating that *H19*-*Igf2* and *Dlk1*-*Dio3* imprinted clusters are the main paternal barriers to normal development of bi-maternal fetus.

Recently, we and others generated mouse androgenetic haploid embryonic stem cells (AG-haESCs) that can be used as “artificial spermatids” to generate alive semi-cloned (SC) mice through intracytoplasmic injection (ICAHCI) into mature oocyte (Li et al., 2012; Yang et al., 2012). We further optimized the SC technology through generation AG-haESCs carrying both *H19*-DMR and *IG*-DMR deletions that can efficiently support the development of SC embryos (Zhong et al., 2015). Interestingly, oocyte-originated haploid ESCs, when removed both *H19*-DMR and *IG*-DMR, could also be used as sperm replacement to produce bi-maternal mice efficiently (Li et al., 2016; Zhong et al., 2016). Importantly, AG-haESCs mediated SC technology combined with CRISPR-Cas9 opens new avenues for genetic analyses *in vivo* (Jiang et al., 2018; Li et al., 2018; Wang and Li, 2019; Wei et al., 2017; Zhong et al., 2015). However, it is still not clear how the *H19*-DMR and *IG*-DMR coordinately regulate SC embryo development.

In this study, we systemically analyzed the roles of paternal *H19*-DMR and *IG*-DMR in SC embryonic development by characterizing the SC embryos at different stages during prenatal development and found that the *H19*-DMR and *IG*-DMR are dispensable for the development of preimplantation of SC embryos. Meanwhile, we showed *H19*-DMR is essential for the development of SC embryos before mid-gestation and *IG*-DMR is critical for late-gestation. Further, we demonstrated that 13-kb deletion of paternal *H19* covering both *H19*-DMR and *H19* gene-body can improve the SC embryonic development before mid-gestation through rescue of placental defects caused by AG-haESCs. Finally, we established AG-haESCs carrying triple deletions (TKO), including *H19*, *H19*-DMR and *IG*-DMR, that can further improve the efficiency of producing viable, normal-sized, and fertile SC mice through ICAHCI.

## RESULTS

### Paternal *H19*-DMR and *IG*-DMR are not essential for the pre-implantation development of SC embryos

In our previous work, we have shown that double deletions of *H19*-DMR and *IG*-DMR (DKO) in mouse AG-haESCs significantly improve the birth rate of SC mice (Zhong et al., 2015). Meanwhile, we found that DKO does not change the transcriptional and methylation profiles of haploid ESCs, suggesting their potential functions during embryonic development. Therefore, in this study, we sought to further define how *H19*-DMR and *IG*-DMR regulate development of SC embryo. To this end, we chose wide-type (WT) AG-haESCs (AGH-OG3) and DKO-AG-haESCs (*H19*^ΔDMR^-*IG*^ΔDMR^-AGH-OG3) carrying an *Oct4-eGFP* transgene, which were derived in our previous studies (Yang et al., 2012; Zhong et al., 2015). PCR and *eGFP* fluorescence activation analyses confirmed that both cells carried expected transgene and deletions (Figures S1A and S1B). The haploidy of AG-haESCs was stably maintained by fluorescence-activated cell sorting (FACS)-based enrichment of haploid cells every 4-6 passages (Figure S1C). We first checked the expression of the imprinted genes regulated by *H19*-DMR and *IG*-DMR through quantitative reverse transcription PCR (RT-qPCR) analysis and found, as expected, *H19* and *Gtl2* were down-regulated in DKO haploid cells, while *Igf2* and *Dio3* were up-regulated (Figure S1D). We then analyzed the transcriptomes of both cells and found that, consistent with previous results (Zhong et al., 2015), WT and DKO haploid cells exhibited a high correlation based on all expressed genes (R = 0.99) (Figure S1E) and all known imprinted genes (R=0.99) (Figure S1F). Interestingly, *Fthl17*, the X-linked maternal imprinted gene that may contribute to the early sex differentiation before gonadal differentiation (Kobayashi et al., 2010), was also downregulated in DKO cells (Figures S1D and S1F). Together, removal of *H19*-DMR and *IG*-DMR in AG-haESCs could mimic the methylation state of *H19*-DMR and *IG*-DMR in sperm that lost during haploid ESC derivation and passaging in 2i medium (Choi et al., 2017; Yagi et al., 2017; Yang et al., 2012) (Figure S1G), leading to DKO haploid ESCs with decreased expression of *H19* and *Gtl2*, two critical paternal imprinted genes for embryonic development.

We next investigated whether *H19*-DMR and *IG*-DMR deletions are benefit to pre-implantation development of SC embryos generated by injection of WT or DKO AG-haESCs into mature oocytes (ICAHCI). The results showed that the efficiency of high-quality blastocysts was similar among sperm carrying *Oct4-eGFP* transgene (control), WT and DKO AG-haESCs (Figures 1A and 1B). To further characterize the similarity of blastocysts produced from different haploid cells, we separated inner cell mass (ICM) and trophectoderm (TE) and carried out low-cell-number (about 10) RNA-seq analysis (Figures S2A-S2C). Hierarchical clustering and principal component analysis (PCA) analyses of transcriptome data showed that all ICM samples were clustered together, but separated from TE samples which were also exclusively clustered together, indicating that the transcriptome differences between all samples are mainly caused by their different lineage origins (Figures 1C and S2D). Pearson correlation analysis further confirmed the similarity among different blastocysts (Figure 1D). Consistently, all imprinted genes exhibited similar expression patterns (Figure 1E). Moreover, the expression levels of *H19*, *Igf2*, *Gtl2* and *Dlk1* were extremely low in all tested ICM and TE samples (Figures 1E and 1F), suggesting that *H19*-*Igf2* and *Dlk1*-*Dio3* imprinted clusters may function during post-implantation development of SC embryos. Therefore, we examined the expression of *H19*, *Igf2*, *Gtl2* and *Dlk1* in control and SC embryos after implantation and found gradually increased expression of four genes following embryonic development from E3.5 to E9.5, implying that DKO could rescue transcriptional defects of *H19* and *Igf2* in SC embryos from WT haploid cells (Figure 1G). These data demonstrate that DKO are not essential for pre-implantation development of SC embryos.

**Figure 1.**
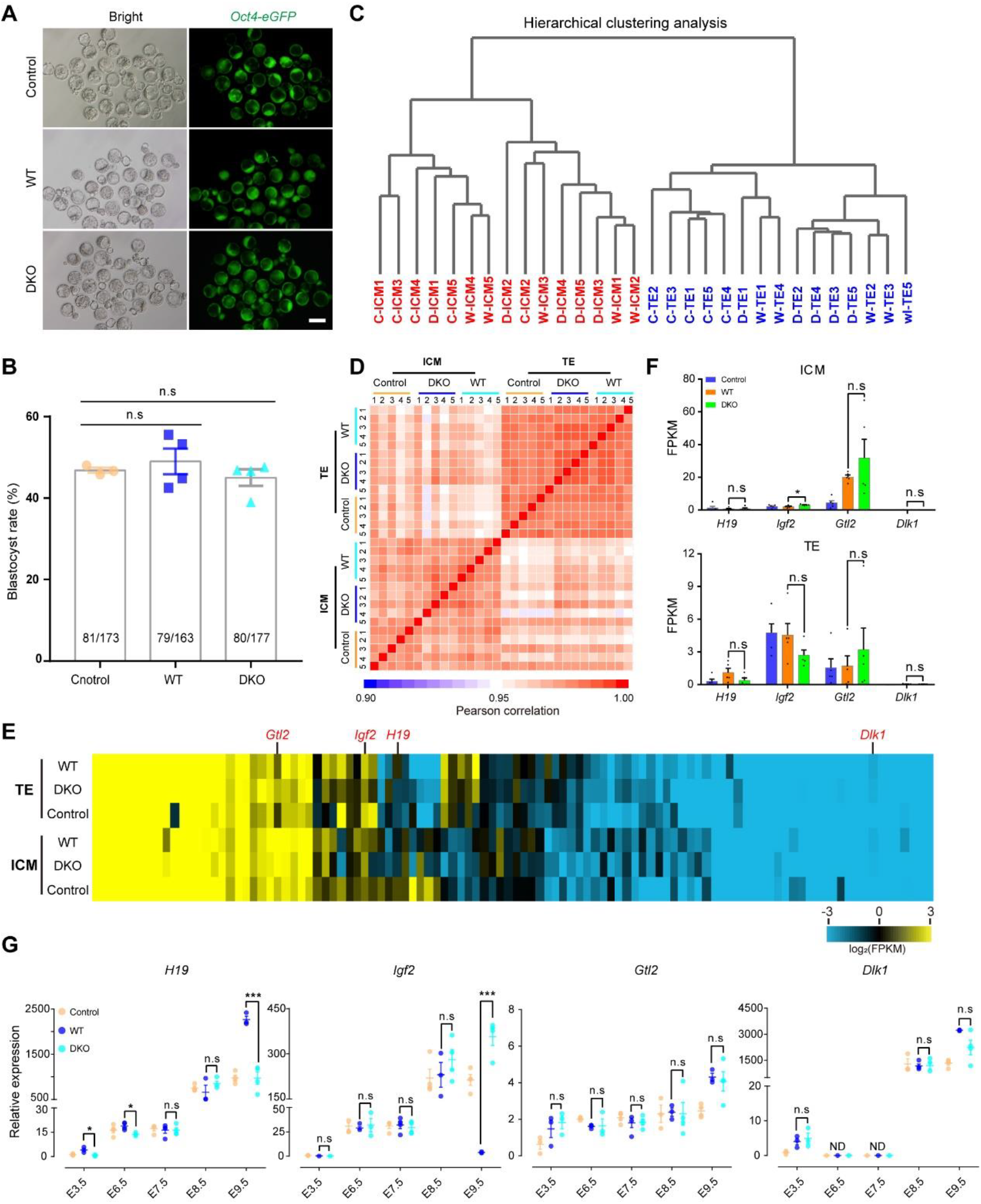
Paternal *H19*-DMR and *IG*-DMR are dispensable for the preimplantation development of SC embryos. (A and B) Representative images of blastocysts (A) and the efficiency of blastocysts derived from sperm (control), WT and DKO-AG-haESCs (B), showing the similar developmental potential among different haploid cells. Numbers in the bars showing blastocysts/two-cells ratio. (C and D) The hierarchical clustering analysis (C) and correlation heatmap (D) of RNA-seq data from ICMs (5 replicates) and TEs (5 replicates) obtained through different haploid donors. C, control embryos derived from intracytoplasmic sperm injection (ICSI); W, embryos from intracytoplasmic WT-AG-haESC injection; D, embryos from DKO-AG-haESC injection. (E) The heatmap showing expression patterns of imprinted genes in TE (top) and ICM (bottom). The gene expression levels were measured as log_2_ (FPKM). (F) The expression levels of *H19*, *Igf2*, *Gtl2* and *Dlk1* in ICM and TE. (G) Transcriptional analysis of *H19*, *Igf2*, *Gtl2* and *Dlk1* genes among control (ICSI), WT and DKO embryos from E3.5 to E9.5 (*n* = 3 embryos). The expression values were normalized to that of *Gapdh*. All error bars in B, F and G indicate the average mean ± s.e.m. \**P* < 0.05, \*\*\**P* < 0.001. n.s, no significant difference.

### *H19* and *IG*-DMR deletions (DKO) significantly improve the developmental potential of SC embryos during mid-gestation

We next examined the effect of DKO on prenatal development by transplantation of 2-cell SC embryos produced through ICAHCI of WT or DKO haploid cells into oviducts of pseudo-pregnant females (Figure 2A). A total of 2173 WT and 1352 DKO SC two-cell embryos were transplanted. Dissection of SC embryos in pregnant females day-by-day from E6.5 to E12.5 revealed that DKO SC embryos implanted at significantly higher efficiency compared with WT SC embryos (41.9% vs 34.6%) (Figure 2B). Meanwhile, the developmental potential of DKO SC was significantly higher than WT SC embryos at E12.5 (Figure 2C). Moreover, higher frequency of abnormal embryos (degeneration or morphological abnormalities) displayed in WT group starting from E8.5 (Figures 2D and 2E). Interestingly, we found that most abnormal DKO SC embryos degenerated by E10.5 and almost all alive DKO SC embryos on E10.5 could develop to term *in vivo* (Figure 2C), whereas a high ratio of WT SC embryos showed both developmental retardation and degeneration and continued to display developmental failure at the following stages (Figures 2F and 2G), suggesting that DKO rescue the developmental defects of SC embryos before mid-gestation.

**Figure 2.**
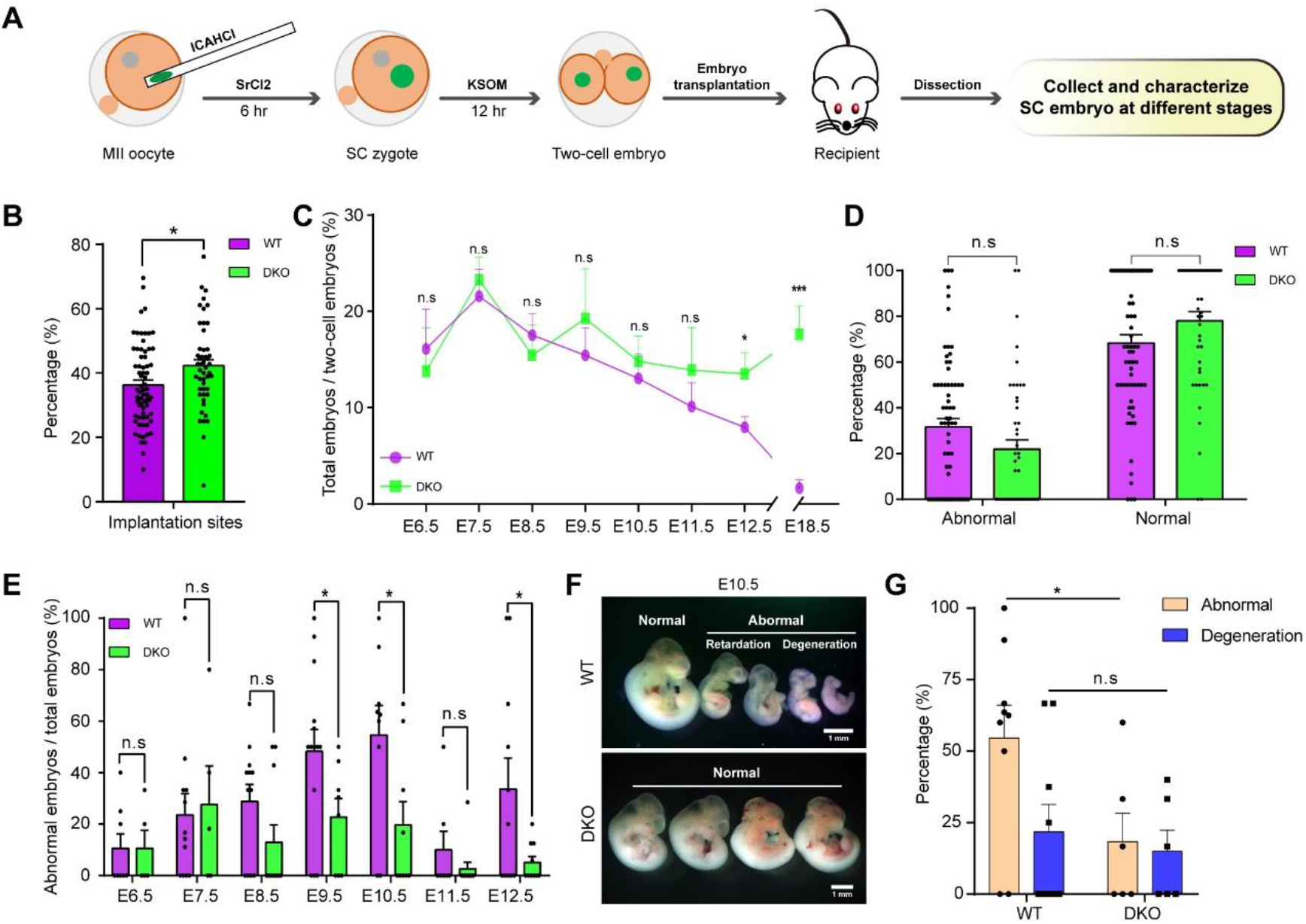
Developmental potential analysis of SC embryos at different stages. (A) Schematic diagram of developmental potential analysis of SC embryos generated via ICHACI of WT or DKO AG-haESCs. (B) Summary of the average implantation rate of WT (*n* = 68 recipients) and DKO (*n* = 49 recipients) SC embryos dissected from E6.5 to E12.5. (C) Comparison of the individual developmental efficiency on different embryonic days between WT and DKO SC embryo. (D) Summary of the total ratios of abnormal and normal SC embryo (from E6.5 to E12.5) derived from WT (*n* = 68 recipients) and DKO (*n* = 49 recipients). (E) Comparison of the individual ratios of abnormal SC embryos on different embryonic days (from E6.5 to E12.5) between WT and DKO groups. (F and G) Representative images of E10.5 SC embryos produced from WT (top panel) and DKO (bottom panel) AG-haESCs (F). The ratios of abnormal and degeneration embryos show in (G). The dark spots in B, D, E and G show one data from one recipient. All error bars in B-E and G indicate the average mean ± s.e.m. \**P* < 0.05, \*\*\**P* < 0.001. n.s means no significant difference.

We next carefully compared the morphological differences between DKO and WT SC embryos. We observed that a high ratio of WT SC fetuses was alive with normal heartbeat on E8.5, but most of them exhibited developmental arrest starting from day 10.5 of gestation (Figure 3A). Particularly, both fetal and placental weights were significantly lower in WT SC conceptus than those of the DKO SC conceptus on E12.5 (Figure 3B). Interestingly, the blood vessel of amniotic sac and umbilical cord were poorer and shorter in the WT SC conceptus, which may influence the nutrient and gas-exchange with mother (Figure 3C). The mature placenta contains three main layers: the labyrinth, which constitutes the main nutrient and gas-exchange surface; the basal zone, which consists of spongiotrophoblast, glycogen cells and different giant cell subtypes; and the maternally derived decidua (Rossant and Cross, 2001). Hematoxylin eosin (HE) staining analysis of placentas indicated that the WT SC placentas had smaller total areas and increased ratio of decidual areas compared with DKO ones (Figures 3D and 3E). Besides, we found that the glycogen cells in the basal zone was dramatically reduced in WT placenta (Figure 3D).

**Figure 3.**
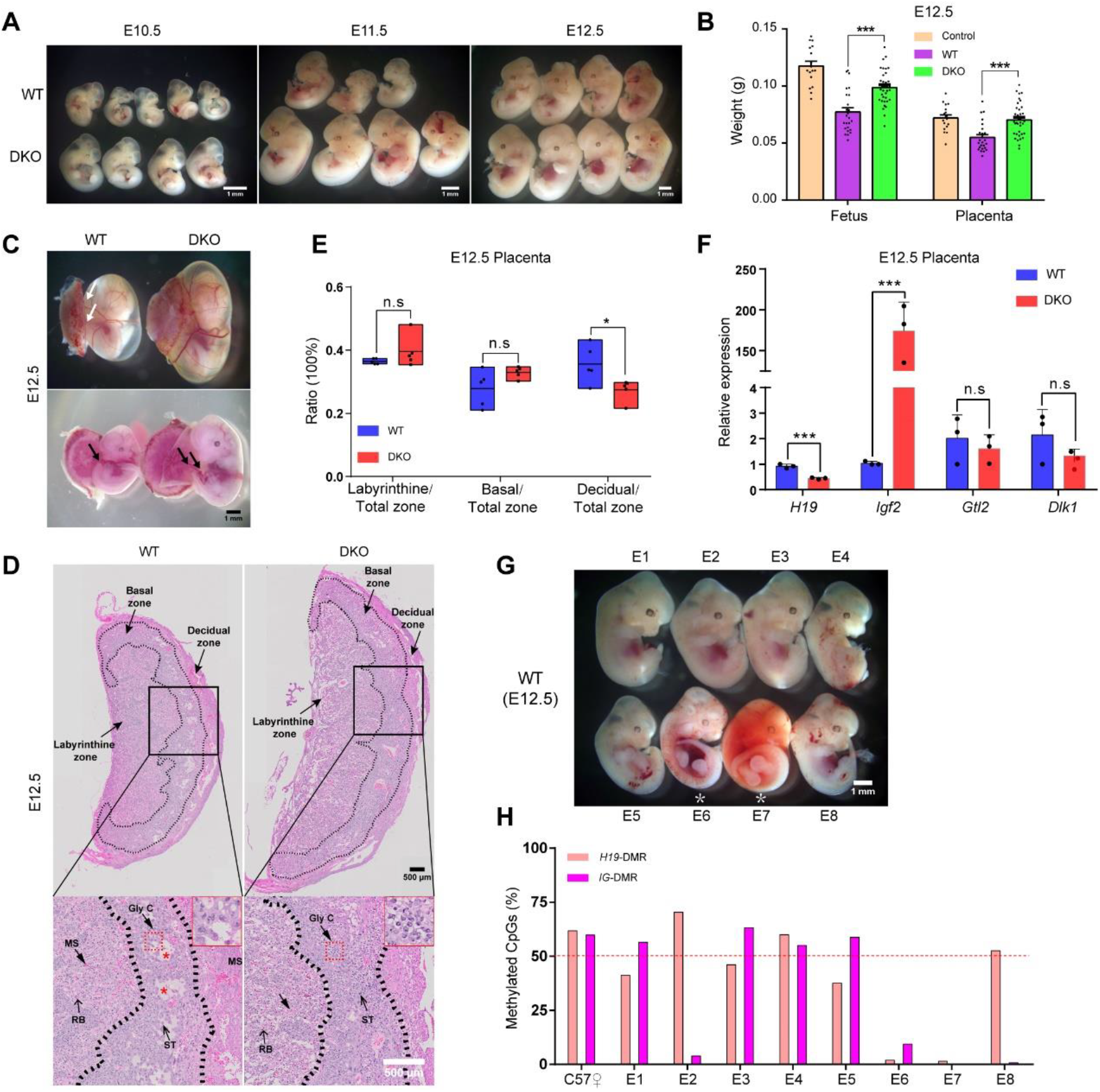
*H19*-DMR and *IG*-DMR double knock-out significantly rescue the retardation of SC embryos at mid-gestation. (A) Representative images of alive SC embryos (E10.5 to E12.5) derived from WT (top panel) and DKO (bottom panel) AG-haESCs show retarded growth in WT group. (B) The fetal and placental weights of E12.5 embryos derived from sperm (control, *n* = 17), WT (*n* = 25) and DKO (*n* = 36) AG-haESCs, showing severe retardation in WT embryos compared with DKO embryos. (C) E12.5 conceptus derived from WT and DKO AG-haESCs. An obvious gap (white arrow) between the placenta and visceral yolk sac shows in WT fetus (top panel). The well-developed umbilical cord (black arrows) shows in the DKO fetus (bottom panel). Experiments were repeated five times independently, with similar results. (D) Whole mount images of SC placentas (E12.5) (top panel). Black boxes in the top panel are magnified in bottom panel. The total placental area is smaller in the WT compared with DKO. Fewer glycogen cells (Gly C) exist in the basal zone of the placenta from the WT compared with DKO, shown in red dashed boxes. Red asterisk showed vacuolation in WT placenta. MS, maternal sinusoid; RB, red blood cells; ST, spongiotrophoblast. Experiments were repeated five times independently, with similar results. (E) The ratios of labyrinthine zone, basal zone and decidual zone to total placental areas of WT and DKO placentas (E12.5), indicating increased decidual area in WT placentas. 5 placentas for each group and at least 5 sections per placenta were analyzed. Middle lines are shown as the mean. (F) Transcriptional analysis of *H19*, *Igf2*, *Gtl2* and *Dlk1* genes in WT and DKO placentas (3 placentas for each group). The expression values were normalized to that of *Gapdh*. Data are shown as the average mean ± s.e.m. in B and F. \**P* < 0.05, \*\*\**P* < 0.001. n.s means no significant difference. (G and H) Images of eight WT SC embryos (G) and their methylation state of the *H19*-DMR and *IG*-DMR (H). Embryos with a white asterisk are dead. The dotted line shows theoretical value of methylation, 50%.

We next assessed the expression of critical imprinted genes in placentas via RT-qPCR. As expected, *H19* was overexpressed whereas *Igf2* expression was significantly decreased in WT placenta (Figure 3F), which may account for the reduced glycogen cells in WT SC placenta (Lopez et al., 1996). *Igf2* is an important regulator in maintenance of the balance between supply and demand systems that is crucial to fine-tune mammalian growth (Angiolini et al., 2011; Constancia et al., 2005). We then tested the expression of glucose transporter genes (*Slc2a1* and *Slc2a3*) and amino acid transporter genes (*Slc38a1*, *Slc38a2*, and *Slc38a4*) in E12.5 placentas. The results showed that *Slc2a3* and *Slc38a2* were significantly increased in the WT SC placenta (Figure S3A), probably leading to increased nutrient transfer that adapts to the nutrient supply to fetal demand (Constancia et al., 2005). Meanwhile, we also found that estrogen receptor (*Esr1* and *Esr2*) and androgen receptor (*Ar*) were highly expressed in WT placenta, whereas the inactivator of testosterone and estrogen (hydroxysteroid 17-beta dehydrogenase 2, *Hsd17b2*) had no difference (Figure S3A), consistent with previous observations that maternal androgen excess reduces placental and fetal weights (Sun et al., 2012). Besides, WT SC fetus died gradually from E11.5 to E15.5 (Figures S3B and S3C) and only two from a total of 308 WT SC (0.6%) derived from AGH-OG3 cells with late passages (over passage 40) survived on E18.5 (Figures S3D and S3E). These results indicate that the developmental failure of WT SC embryos is mainly caused by placental defects, which can be rescued by *H19*-DMR and *IG*-DMR deletions.

### *IG*-DMR deletion alone fails to rescue the developmental retardation of SC embryos before E12.5

Interestingly, we observed that all dead embryos (E12.5) from WT AG-haESCs lost DNA methylation at the *H19*-DMR (Figures 3G and 3H). In contrast, loss of DNA methylation at the *IG*-DMR appeared in both live and dead embryos on E12.5 (Figures 3G and 3H). These results suggest that *H19*-DMR methylation is essential for SC embryonic development after implantation, while *IG*-DMR may not be very important before mid-gestation. To further explore their roles in SC embryonic development, we firstly deleted the 4kb region of *IG*-DMR (Lin et al., 2003) in AGH-OG3 cells using CRISPR/Cas9 technology (Figure 4A). Two single guide RNAs (sgRNAs) targeting upstream and downstream of the region were designed and ligated to pX330-*mCherry* plasmid (Wu et al., 2013) and transfected into haploid cells, leading to two stable cell lines (termed *IG*^ΔDMR^-AGH-OG3-1 and 2) (Figure S4A). Interestingly, these two cell lines were free of DNA methylation at *H19*-DMR (Figure S4B). ICAHCI analysis indicated that *IG-*DMR deletion did not improve developmental potential of SC embryos, and the percentage of abnormal SC embryos was comparable to that of WT SC embryos (Figure 4B). Meanwhile, the alive E12.5 fetus of *IG*^ΔDMR^ exhibited growth-retarded phenotypes in both embryos and placentas (Figures 4C and 4D). Further, placental defects were more severe than WT fetus (Figures 4D-4F and S4C). Consistently, DNA methylation at *H19*-DMR was absent in *IG*^ΔDMR^ SC embryos (Figures S4D and S4E), resulting in misexpression of *H19* and *Igf2* in *IG*^ΔDMR^ SC placentas (Figure 4G). As expected, SC embryos with paternal *IG*-DMR deletion alone rarely developed to day 18.5 gestation (Figure 4H). These results indicate that *IG*-DMR deletion in AG-haESCs is dispensable for the early stages of SC embryo development.

**Figure 4.**
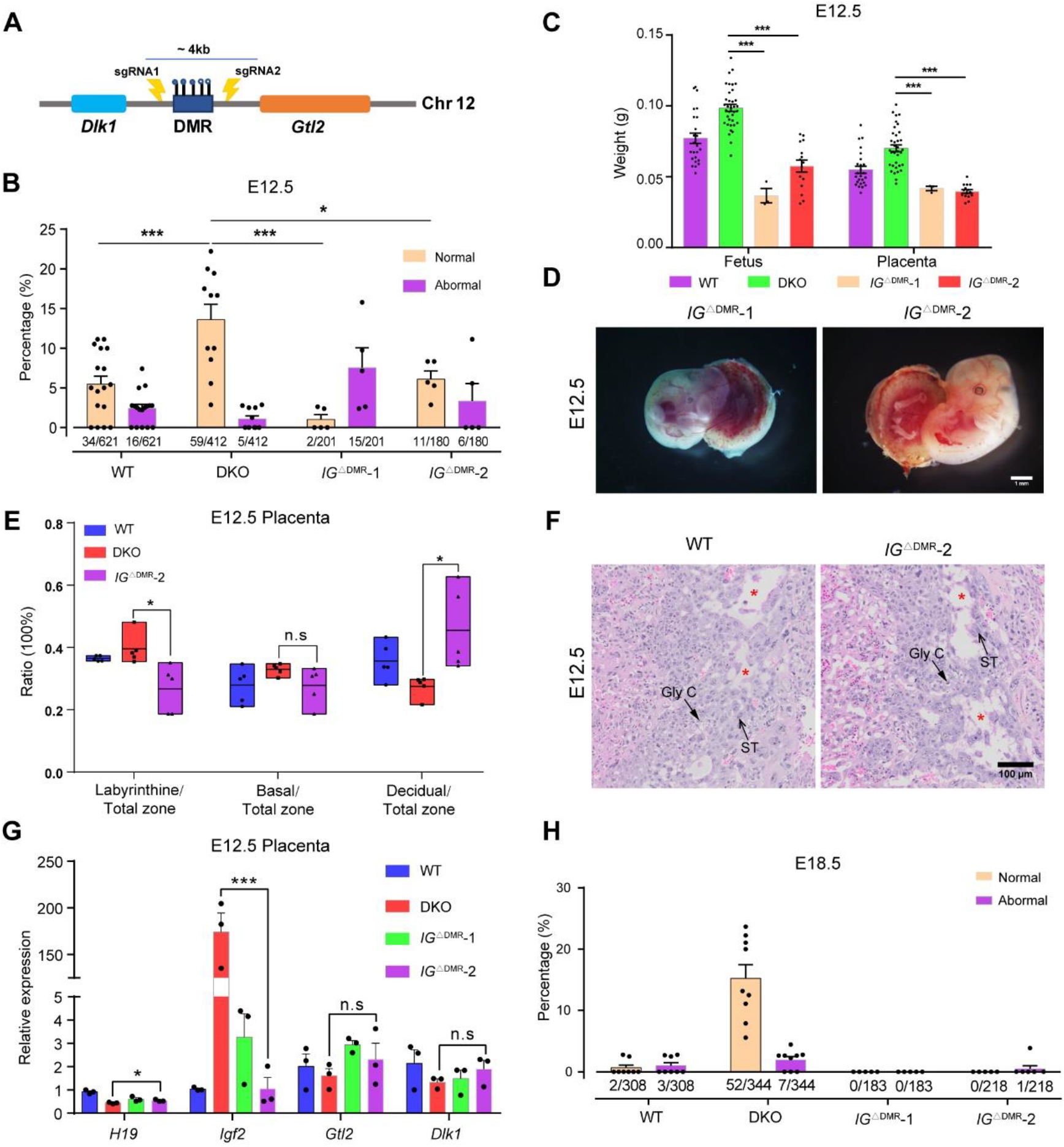
Deletion of *IG*-DMR alone fails to rescue the developmental retardation of SC embryos before E12.5. (A) Schematic diagram of sgRNAs targeting for the removal of *IG*-DMR. A dark blue bar represents the deleted region (about 4-kb). (B) The ratios of normal and abnormal SC embryos (E12.5) in transferred 2-cell embryos derived from WT (*n* = 17 recipients), DKO (*n* = 11 recipients) and *IG*^ΔDMR^ (*n* = 5 recipients for each cell line) AG-haESCs. Numbers under the bars indicate E12.5 embryos/transferred two-cell embryos. (C) The fetal and placental weights of E12.5 embryos derived from WT (*n* = 25), DKO (*n* = 36) and *IG*^ΔDMR^ (*n* = 5 for *IG*^ΔDMR^-1 and *n* = 15 for *IG*^ΔDMR^-2) AG-haESCs, showing severe retardation in *IG*^ΔDMR^ SC embryos compared with DKO. (D) Severe developmental retardation shown in *IG*^ΔDMR^ SC fetuses (E12.5). Experiments were repeated five times independently, with similar results. (E) The ratios of labyrinthine zone, basal zone and decidual zone to total placenta areas of WT, DKO and *IG*^ΔDMR^ placentas (E12.5), indicating decreased labyrinthine zone and increased decidual area in *IG*^ΔDMR^ placentas compared with DKO, *n* = 5 placentas for each group and at least 5 sections per placenta analyzed. Middle lines are shown as the mean. (F) Histological sections of the placentas from WT and *IG*^ΔDMR^ SC fetus on E12.5. Red asterisks show vacuolation in WT and *IG*^ΔDMR^ placentas. Gly C, glycogen cells; ST, spongiotrophoblast. Experiments were repeated five times independently, with similar results. (G) Transcriptional analysis of *H19*, *Igf2*, *Gtl2* and *Dlk1* genes in WT, DKO and *IG*^ΔDMR^ placentas (*n* = 3 placentas for each group). The expression values were normalized to that of *Gapdh*. (H) The ratios of normal and abnormal SC embryos (E18.5) in transferred 2-cell embryos derived from WT (*n* = 8 recipients), DKO (*n* = 9 recipients) and *IG*^ΔDMR^ (*n* = 5 recipients for each cell lines) AG-haESCs. Numbers under the bars indicate E18.5 embryos/transferred two-cell embryos. Data are shown as the average mean ± s.e.m in B, C, G and H. \**P* < 0.05, \*\*\**P* < 0.001. n.s means no significant difference.

### *H19*^Δ13kb^ rescues developmental defects of SC embryos at mid-gestation

Having demonstrated that *IG*-DMR is not critical for SC embryonic development before mid-gestation, we next examined the role of *H19*-DMR in SC embryonic development. Due to that *H19*-DMR deletion cannot totally repress paternal *H19* expression and results in reduced *Igf2* expression (Thorvaldsen et al., 1998; Thorvaldsen et al., 2002), to exclude the influence of paternal *H19* expression on the development of SC embryos, we attempted to remove the 13-kb region of *H19* that includes the gene body and the DMR in AG-haESCs (AGH-OG3) (Figures 5A and S5A). Two stable cell lines (*H19*^Δ13kb^-AGH-OG3-1 and 2) were generated and harbored an abnormal methylation status at *IG*-DMR similar to AGH-OG3 cells (Figures S5B and S5C). ICAHCI analysis showed that *H19*^Δ13kb^ led to comparable developmental potential of SC to DKO AG-haESCs (Figure 5B) at E12.5, probably through rescue of placental defects (Figures 5C-5F, S5D and S5E). Consistently, the expression of imprinted genes in *H19*^Δ13kb^ placentas were similar to those in *H19*^ΔDMR^-*IG*^ΔDMR^ DKO placentas (Figure 5G). Interestingly, we observed fewer abnormal embryos in *H19*^Δ13kb^ group compared to DKO group, probably due to slightly higher level of *Igf2* expression in *H19*^Δ13kb^ embryos (Figure 5G).

**Figure 5.**
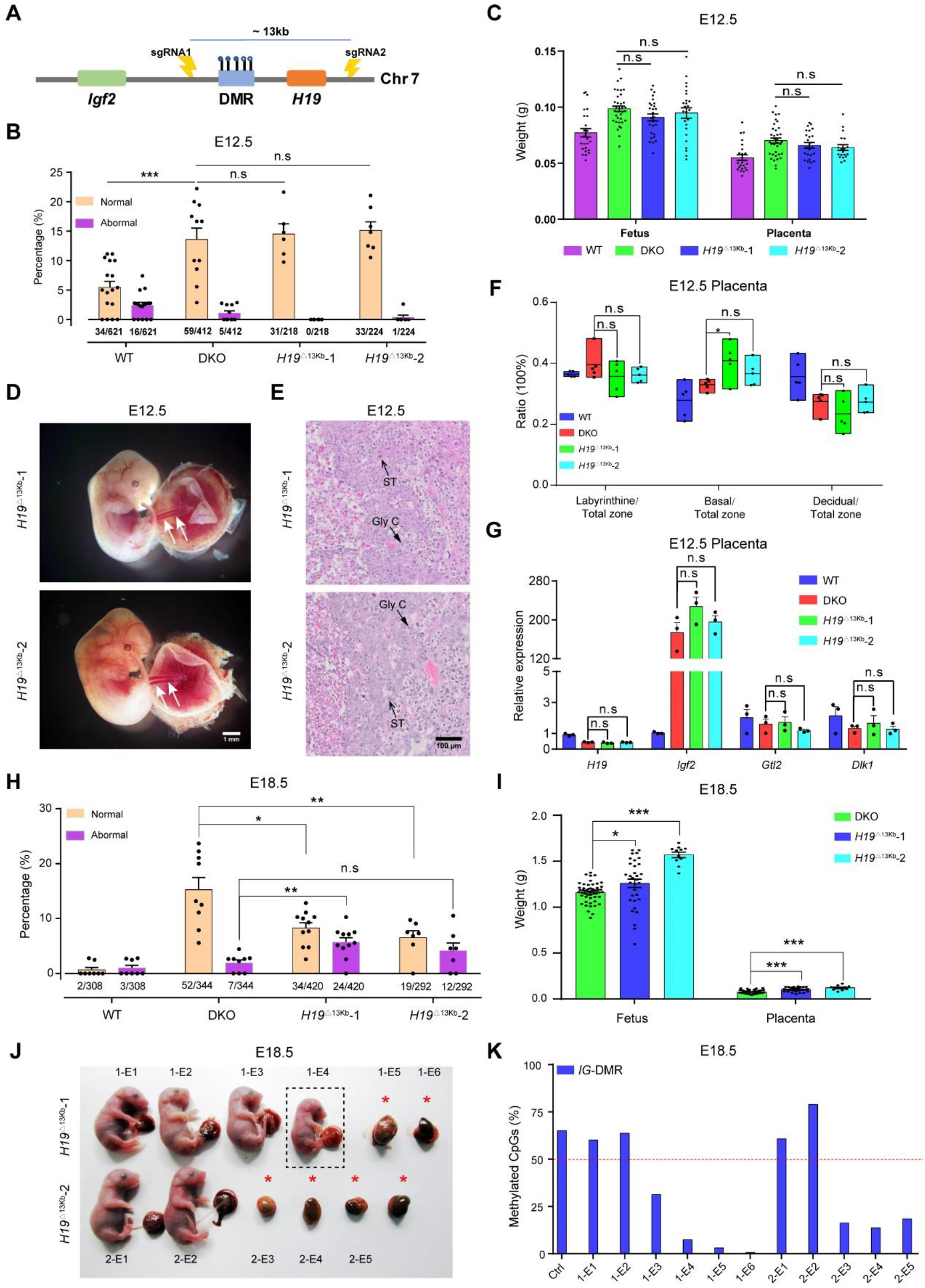
*H19*^Δ13kb^ rescues the abnormal development of SC embryos at mid-gestation. (A) Schematic diagram of sgRNAs targeting for the removal of *H19*-DMR and *H19* transcription unit (about 13-kb). (B) The ratios of E12.5 normal and abnormal SC embryos in transferred 2-cell embryos derived from WT (*n* = 17 recipients), DKO (*n* = 11 recipients) and *H19*^Δ13kb^ (*n* = 6 recipients for *H19*^Δ13kb^-1 and *n* = 7 for *H19*^Δ13kb^-2) AG-haESCs. Numbers under the bars indicate E12.5 embryos/transferred two-cell embryos. (C) The fetal and placental weights of E12.5 SC embryos derived from WT (*n* = 25), DKO (*n* = 36) and *H19*^Δ13kb^ (*n* = 27 for cell line1 and *n* = 25 for cell line2) AG-haESCs, showing comparable developmental potential of *H19*^Δ13kb^ and DKO SC embryos. (D) Images of *H19*^Δ13kb^ SC fetuses (E12.5) showing normal development. White arrows show the umbilical cords. Experiments were repeated five times independently, with similar results. (E) Histological sections of the placentas from *H19*^Δ13kb^ SC fetus at day 12.5 gestation. Gly C, glycogen cells; ST, spongiotrophoblast. Experiments were repeated five times independently, with similar results. (F) The ratios of labyrinthine zone, basal zone and decidual zone to total placenta areas of WT, DKO and *H19*^Δ13kb^ placentas (E12.5), indicating comparable development of DKO and *H19*^Δ13kb^ placentas. *n* = 5 placentas for each group and at least 5 sections per placenta were analyzed. Middle lines are shown as the mean. (G) Transcriptional analysis of *H19*, *Igf2*, *Gtl2* and *Dlk1* genes in WT, DKO and *H19*^Δ13kb^ placentas (*n* = 3 placentas for each group). The expression values were normalized to that of *Gapdh*. (H) The ratios of normal and abnormal SC embryos (E18.5) in transferred 2-cell embryos derived from WT (*n* = 8 recipients), DKO (*n* = 9 recipients) and *H19*^Δ13kb^ (*n* = 11 recipients for *H19*^Δ13kb^-1 and *n* = 7 for *H19*^Δ13kb^-2) AG-haESCs. Numbers under the bars indicate E18.5 embryos/transferred two-cell embryos. (I) The fetal and placental weights of E18.5 SC embryos derived from DKO (*n* = 51) and *H19*^Δ13kb^ (*n* = 32 for *H19*^Δ13kb^-1 and *n* = 11 for *H19*^Δ13kb^-2) AG-haESCs, showing highly developmental potential of *H19*^Δ13kb^ compared to DKO SC embryos. Data are shown as the average mean ± s.e.m in B, C and G-I. \**P* < 0.05, \*\**P* < 0.01, \*\*\**P* < 0.001. n.s means no significant difference. (J and K) Images of *H19*^Δ13kb^ SC embryos on E18.5 (J) and their methylation state in the *IG*-DMR (K) The black dotted box shows the retard embryo and red asterisks show degenerative embryos. The dotted line in (K) shows theoretical value of methylation (50%).

We next checked the developmental potential of *H19*^Δ13kb^ SC fetus at late-gestation. Compared to DKO SC embryos, the percentage of *H19*^Δ13kb^ SC embryos which normally developed to E18.5 was significantly reduced, accompanied with higher frequency of abnormal development (Figure 5H). As expected, the survived *H19*^Δ13kb^ SC embryos showed higher fetal and placental weights compared to DKO embryos (Figure 5I). Notably, we observed that the DNA methylation at *IG*-DMR lost in all abnormal *H19*^Δ13kb^ embryos on E18.5 (Figures 5J and 5K). Furthermore, growth-retarded *H19*^Δ13kb^ SC embryos with low methylation of *IG*-DMR (Figures S6A and S6B) exhibited enhanced *Gtl2* expression but repressed *Dlk1* and *Dio3* expression in different tissues (Figure S6C), indicating the critical roles of *Dlk1*-*Dio3* imprinted cluster in late-gestation development.

### *H19*^Δ13kb^ rescues developmental defects of SC embryos at mid-gestation through rescue of placental defects

Since different deletions resulted in different phenotypes in placentas at E12.5, we then sought to explore whether deletions changed the gene expression in the placenta by performing RNA sequencing (RNA-seq) analysis. To do this, we collected E12.5 placentas of five groups (three replicates for each group) derived from sperm (ICSI), WT AG-haESCs (WT), *H19*^ΔDMR^-*IG*^ΔDMR^-AG-haESCs (DKO), *IG*^ΔDMR^-AG-haESCs (*IG*^ΔDMR^) and *H19*^Δ13kb^-AG-haESCs (*H19*^Δ13kb^) respectively. The results indicated that WT and *IG*^ΔDMR^ placentas were clearly clustered together, and exhibited transcriptome patterns different from ICSI, *H19*^Δ13kb^ and DKO placentas (Figures 6A, S7A and S7B). These results were consistent with the observations that *H19*^Δ13kb^ and DKO significantly rescued the placental defects, while *IG*^ΔDMR^ showed severe placenta defects as WT (Figures 4C-4F and 5C-5F). We further compared the expression patterns of imprinted genes and found that two groups of imprinted genes were differentially expressed between DKO/*H19*^Δ13kb^/ICSI and *IG*^ΔDMR^/WT, including genes involved in *H19*-*Igf2* and *Dlk1*-*Dio3* imprinted clusters (Figures 6B and 6C). Moreover, we also found three classes of genes which showed differential expression levels among the five groups of placentas (Figure 6D). Class I (*n* = 64) showed lower expression in *IG*^ΔDMR^/WT compared with DKO/*H19*^Δ13kb^/ICSI and was associated with cell cycle; Class II (*n* = 137) showed higher expression in *IG*^ΔDMR^/WT and was enriched for developmental genes; Class III (*n* = 55) only expressed in WT and is associated with chromatin assembly (Figures 6D and 6E). Interestingly, we noticed that some *Hox* genes were abnormally activated in *IG*^ΔDMR^/WT (Figure 6F), implying that they may be involved in abnormal SC embryonic development. A previous study has shown that *H19* and *Igf2* maintain inactive during the first cell fate decision (ICM and TE formation), while they are specifically activated in visceral endoderm (VE) cells that mainly contribute to placental development, but not in epiblast (Epi) cells that develop into embryonic tissues (Figure 6G) (Zhang et al., 2018). These observations suggest that *H19* and *H19*-DMR may play a role in repression of development-related genes in placenta. In summary, *IG*^ΔDMR^ cannot rescue the abnormal gene expression in WT, including imprinted genes and other functional genes in E12.5 placentas, which may lead to severe developmental defects of SC embryos; in contrast, DKO and *H19*^Δ13kb^ correct the misexpression of imprinted genes and other important functional genes in placenta, which contributes to the normal development in placenta. Taken together, these results provide strong evidence to demonstrate that *H19-Igf2* locus on the paternal genome plays a key role in the SC embryonic development before mid-gestation, whereas *Dlk1*-*Dio3* locus on the paternal genome is indispensable for normal embryonic development of late-gestation.

**Figure 6.**
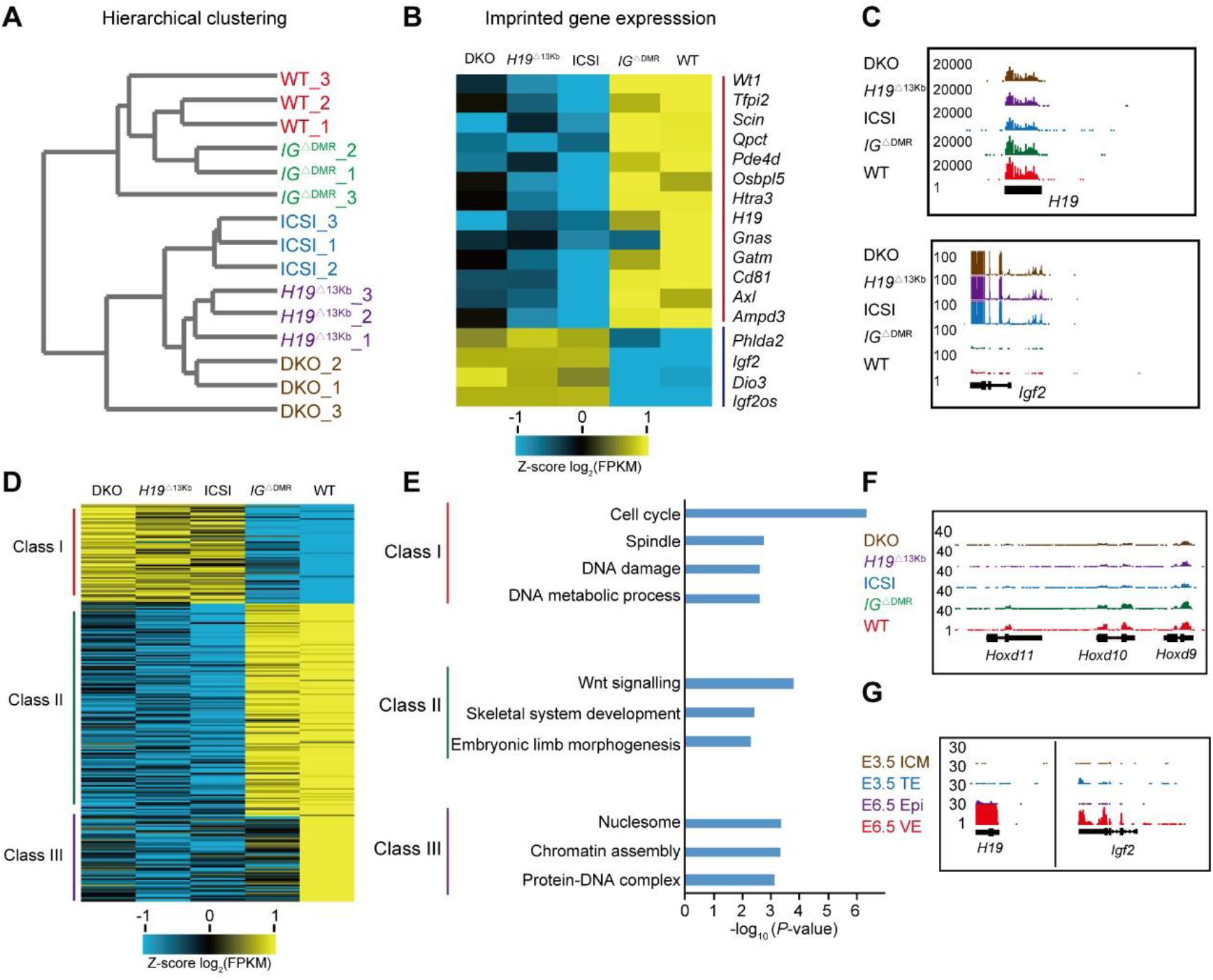
Global transcriptome analysis of SC placentas with different deletions. (A) The hierarchical clustering analysis of RNA dataset from placentas obtained through oocyte injection of sperm, WT, *IG*^ΔDMR^, *H19*^Δ13kb^ and DKO AG-haESCs respectively. (B) Differentially expressed imprinted genes in different placentas. Representative genes enriched in each cluster are shown in the right column. (C) UCSC Genome Browser (University of California, Santa Cruz) snapshots of *H19* and *Igf2* expression levels in different placentas. (D and E) The heatmap showing 3 classes of differentially expressed genes among different placentas (D) and Gene Ontology (GO) analysis of genes in the 3 classes (E). (F) Genome Browser snapshots of *Hoxd11*, *Hoxd10* and *Hoxd9* expression levels in different placentas. (G) Genome Browser snapshots of *H19* and *Igf2* expression levels in E3.5 and E6.5 embryos, according to previous data (Zhang et al., 2018).

### Triple deletions of *H19*, *H19*-DMR and *IG*-DMR further improve the developmental potential of SC embryos

Given that *H19*^Δ13kb^ can give rise to better development compared to DKO group on E12.5 (Figure 5B), we asked whether combined deletions between *H19*^Δ13kb^ (including *H19* and *H19*-DMR) and *IG*-DMR (TKO-AG-haESCs) could further improve full-term developmental potential of SC embryos. Two stable cell lines were generated through removal of *IG*-DMR in *H19*^Δ13kb^-AGH-OG3-1 (termed *H19*^Δ13kb^-*IG*^ΔDMR^-AGH-OG3-1 and 2 or TKO-1 and 2) and injected into oocytes to produce mice (Figures 7A, 7B, S8A and S8B). The results showed that TKO-1 exhibited comparable developmental efficiency at E18.5 to DKO cells, while TKO-2 cells displayed a significantly higher efficiency (Figure 7B). Interestingly, TKO SC embryos exhibited remarkably higher body and placenta weights than DKO fetus, reaching to the levels of ICSI pups (Figures 7C and 7D). These could be caused by total deletion of paternal *H19*, resulting in decreased *H19* expression in major organs during embryonic development (Figure 7E).

**Figure 7.**
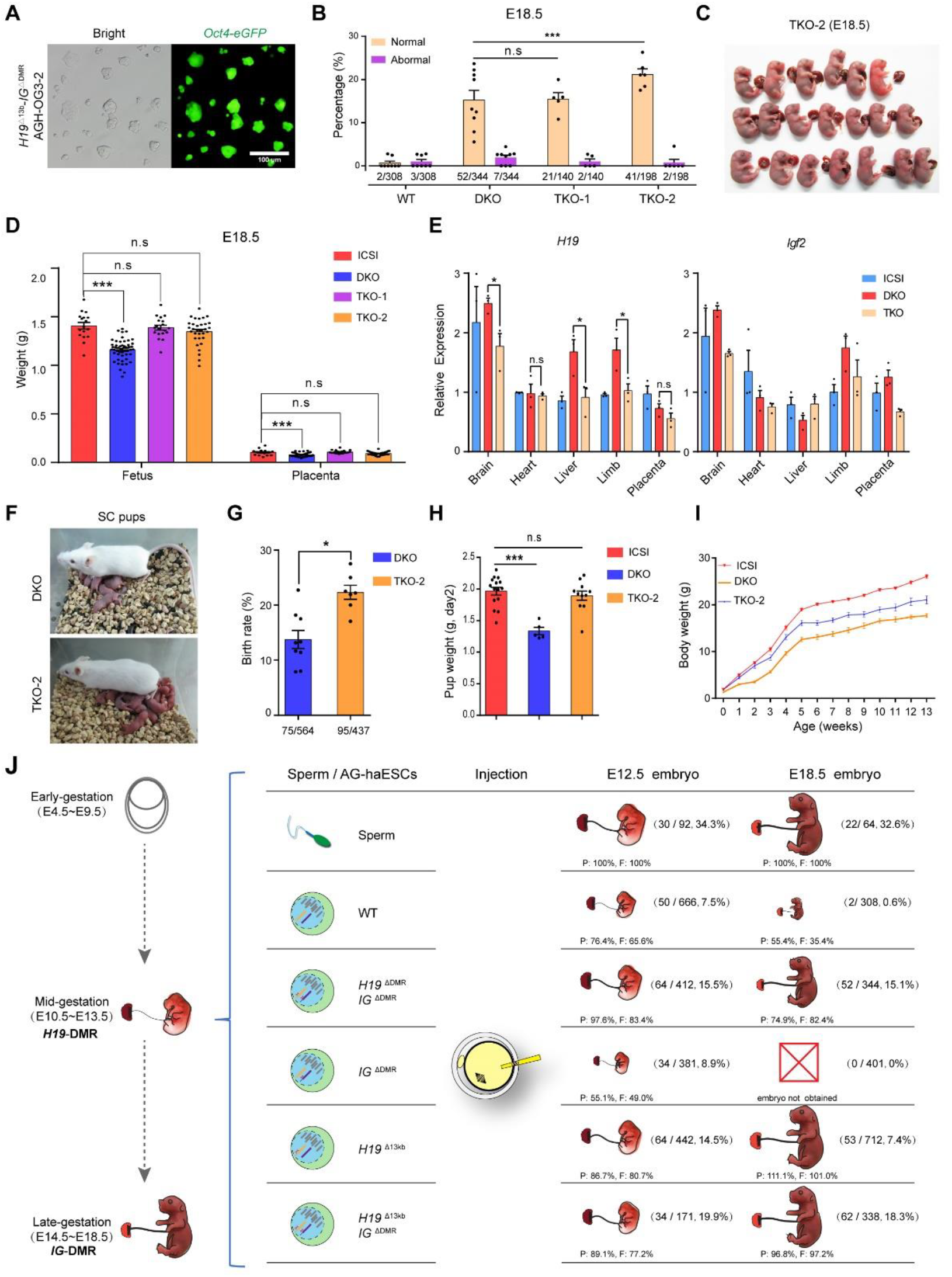
Combined deletions of *H19*, *H19*-DMR and *IG*-DMR in AG-haESCs further increases the prenatal and postnatal development of SC embryos. (A)Images of a TKO-AG-haESC (*H19*^Δ13kb^-*IG*^ΔDMR^-AGH-OG3-2) line carrying an *Oct4-eGFP* transgene. (B)The ratios of normal and abnormal E18.5 SC embryos in transferred 2-cell SC embryos derived from WT (*n* = 8 recipients), DKO (*n* = 9 recipients) and TKO (*n* = 5 recipients for TKO-1 and 6 for TKO-2) AG-haESCs. Numbers under the bars indicate E18.5 embryos/transferred two-cell embryos. (C) Images of E18.5 TKO SC embryos derived from TKO-2 cells. Experiments were repeated five times independently, with similar results. (D) The fetal and placental weights of E18.5 embryos derived from sperm (*n* = 17), DKO (*n* = 51) and TKO (*n* = 19 for TKO-1 and *n* = 32 for TKO-2) AG-haESCs, showing comparable development of TKO SC and control embryos (ICSI). (E) Transcriptional analysis of imprinted genes *H19* and *Igf2* in different tissues of E18.5 fetuses (ICSI, DKO and TKO), including brain, heart, liver, limb and placenta (*n* = 3 samples for each group). The expression values were normalized to that of *Gapdh*. (F and G) The birth rates of SC pups from ICAHCI experiments using DKO and TKO-2 cells (G). Representative images of recipient mouse and SC pups shown in (F). (H) Higher body weight of TKO-2 SC pups than that DKO pups on day 2 after birth. (I) Growth curve of ICSI mice (females, *n* = 9), DKO (*n* = 4) and TKO-2 SC mice (*n* = 10), showing that TKO further improve the postnatal developmental potential of SC mice. Data are shown as the average mean ± s.e.m in B, D, E and G-I. \**P* < 0.05, \*\*\**P* < 0.001. n.s means no significant difference. (J) Summary of the temporal regulation of paternal genomic imprinting (*H19*-DMR and *IG*-DMR) during SC embryonic development. The percentages of placental (P) and fetal (F) weights are based on the placental and fetal weights of ICSI. The numerator in parentheses means the alive embryos obtained and denominator means the total number of transferred two-cell embryos.

Importantly, the birth rate and body weight of TKO-2 newborn pups were notably higher than DKO (Figures 7F-7H). Furthermore, the growth profiling analysis showed that TKO-2 enhanced the growth of SC mice (Figure 7I). Consistently, removal of *H19* gene-body in another DKO cell line (termed *H19*^ΔDMR^-*IG*^ΔDMR^-AGH-2 or O48) generated previously (Zhong et al., 2015) resulted in a stable cell line termed O48-TKO, from which, SC pups with higher birth weight were produced (Figures S8C-S8E). Finally, we tested the reproductive ability of TKO SC mice by collecting mature oocytes from superovulated 4-weeks-old SC mice. The average number of oocytes from each mouse was similar among TKO, DKO and ICSI groups (Figures S8F and S8G). Besides, the concentration of luteinizing hormone (LH) in plasma of adult mice (4-month old) were also comparable between SC and control mice (Figure S8H). Moreover, TKO mice grew to adulthood and reproduced normally (Figure S8I). Genotyping analysis of a total of 9 live-born pups from 2 litters of progeny delivered by TKO SC mice showed that 3 carried the 13kb-deletion of *H19* (Figure S8J). All pups harboring maternal deletion of *IG*-DMR died shortly after birth, consistent with previous reports that maternal transmission of the *IG*-DMR deletion causes postnatal or neonatal lethality (Lin et al., 2003; Zhong et al., 2015). Taken together, these results demonstrate that TKO can further improve prenatal and postnatal development of SC embryos.

## DISCUSSION

Previous studies have shown that ng oocytes carrying both *H19* and *IG*-DMRs can efficiently support bi-maternal embryonic development after fusion with a mature oocyte (Kawahara et al., 2007). Meanwhile, we showed that removal of these two DMRs in AG and PG-haESCs can results in high-efficiency generation of SC and bi-maternal mice respectively through injection into oocytes (Zhong et al., 2016; Zhong et al., 2015). Moreover, we found that *H19* and *IG*-DMR deletions do not change the transcriptional and methylation profiles in haESCs (Zhong et al., 2016; Zhong et al., 2015), implying that *H19* and *IG*-DMRs may function during embryonic development. However, it is still largely unknown how *H19* and *IG*-DMRs coordinately regulate SC embryonic development. In this study, we show that *H19*-DMR and *IG*-DMR temporally regulate SC embryo development (Figure 7J). *H19*-DMR and *IG*-DMR are dispensable for the preimplantation development of SC embryos. *H19*-DMR is indispensable for mid-gestation development, while *IG*-DMR plays a critical role in late-gestation development. *H19*-DMR is important for placental development before mid-gestation and double deletions of *H19* and *H19*-DMR further improve SC embryonic development at mid-gestation, probably through regulation of *Igf2* expression. Importantly, triple knockout of *H19*, *H19*-DMR and *IG*-DMR further improve the prenatal and postnatal development of SC mice.

Mice carrying paternal *H19*-DMR or *IG*-DMR deletion that partially mimics methylation state of DMR (Barlow and Bartolomei, 2014) have been generated a decade ago (Lin et al., 2003; Thorvaldsen et al., 2002), displaying minimal if any adverse phenotypes. Therefore, it is expectable that sperm with both *H19*-DMR and *IG*-DMR deletions obtained through multiple rounds of breeding between two mutant mouse lines may also result in overall normal embryonic development, thus impeding the study of coordinated regulation between two imprinted loci in embryonic development. In contrast, AG-haESCs with low methylation level especially in imprinted regions results in abnormal embryonic development upon injection into oocytes (Yang et al., 2012). Interestingly, after removal of *H19*-DMR and *IG*-DMR in AG-haESCs to rescue the methylation lost at DMRs, the resultant DKO-AG-haESCs can efficiently support normal SC embryonic development, leading to alive pups with a high birth rate comparable to that of round spermatid injection (ROSI) (Zhong et al., 2015). Taken together, these results suggest that AG-haESCs, acting as the sperm replacement, can be used to study the imprinting function *in vivo*. Using this technology, our current study reveals the temporal regulation of paternal *H19*-*Igf2* and *Dlk1*-*Dio3* during embryonic development, providing a strong evidence to demonstrate AG-haESCs as an unique tool to study genomic imprinting *in vivo* (Li and Li, 2019).

Due to the accessibility of genomic modifications in AG-haESCs, this technology may be extended to different topics of imprinting studies *in vivo*, such as: 1) identification of maternal imprinted genes that are critical for embryonic development, whose deletion in AG-haESCs may result in SC embryonic lethal; 2) investigation of control elements of the imprinted locus through one-step generation of SC mice carrying different modifications in the imprinted locus by respective injection of AG-haESCs with corresponding mutations into oocytes; 3) characterization of the expression pattern of a imprinted gene during embryonic development through insertion of a standard tag endogenously in AG-haESCs; and 4) interaction analysis between different imprinted genes/loci.

TKO-AG-haESCs generated in this study exhibit higher developmental potential and better postnatal development of SC embryos; these cells, combing with CRISPR-Cas9 technology, may further enhance genetic analyses *in vivo* (Li et al., 2018; Wang and Li, 2019; Wei et al., 2017). Further investigation into *H19-Igf2*, *Dlk1-Dio3* or other loci will not only reveal mechanisms involved in embryonic development, but may also help to fine-tune the embryonic development for more efficient generation of SC mice.

## MATERIALS AND METHODS

### Animals

All mice were housed in individual ventilated cages (IVC) under specific-pathogen-free and a 12:12 h light/dark cycle condition. MII oocytes are all from B6D2F1 (C57BL/6♀× DBA/2♂) female mice. The *Oct4-eGFP* males (C57BL/6 background) provided sperm for ICSI experiments. All the pseudopregnant foster mothers were ICR females. All animal procedures were performed under the ethical guidelines of the Shanghai Institute of Biochemistry and Cell Biology, Chinese Academy of Sciences, Shanghai, China.

### Plasmids construction

To generate CRISPR-Cas9 plasmid for gene manipulation, specific sgRNAs were synthesized (Sunya, Shanghai), annealed, and fused to BpiI (Thermo Scientific™) linearized pX330-*mCherry* plasmid (Addgene#98750). All inserted sgRNA sequences were validated by Sanger sequencing.

### Cell culture and transfection

Mouse androgenetic haploid embryonic stem cells (AG-haESCs) of AGH-OG3 and *H19*^ΔDMR^*-IG*^ΔDMR^-AGH-OG3 were from our previous works(Yang et al., 2012; Zhong et al., 2015), respectively. These cells were cultured in DMEM (Merk) with 15% FBS (Thermo Scientific™), Penicillin-Streptomycin, Non Essential Amino Acids, Nucleosides, L-Glutamine Solution, 2-Mercaptoethanol, 1000 U/ml Lif, 1 μM PD03259010 (Selleck) and 3 μM CHIR99021 (Selleck). Cells were transfected using Lipofectamine 3000 reagent (Thermo Scientific™) according to the manufacturer’s protocols.

### CRISPR/Cas9 mediated imprinted region deletion

For constructing gene-edited AG-haESCs, cells were transfected with corresponding pX330-*mCherry* plasmids, including specific sgRNAs (Table S1). 24 hours after transfection, the haploid cells expressing red fluorescence protein were enriched with flow cytometry (FACS AriaII, BD Biosciences) and plated at low density (about 8000 cells per well of 6-well plate). One week after plating, single colony was picked for derivation of gene manipulation AG-haESCs identified by PCR and Sanger sequencing. For the enrichment of haploid cells, cells were trypsinized, incubated with 15 μg/ml Hoechst 33342 (Thermo Scientific™) in 37 °C water bath for 5-10 min, transferred through a 40-μm cell strainer and enriched by sorting 1N peak on AriaII.

### Genomic DNA extraction and genotyping

Mouse tails and AG-haESCs were lysed by Mouse Direct PCR Kit (Bimake) according to the manufacturer’s guidance. After centrifuge, 12,000 rpm for 10 min, 1-2 μl supernatant of digested solution can be used as the PCR template. The genome extraction from mouse embryos or pups were using the TIANamp Genomic DNA Kit (TIANGEN), which used for genotyping or bisulphite sequencing.

### Bisulphite sequencing

The bisulfite conversion was performed into one-step using the EZ DNA Methylation-GoldTM Kit (ZYMO research) for 500 ng genomic DNA, following the manufacturer’s instructions. The recovery DNA products were amplified by a first round of nested PCR, following a second round using loci specific PCR primers (Table S1). The amplified products were purified by gel electrophoresis using Universal DNA Purification Kit (TIANGEN) and cloned into a pMD19-T vector (Takara). For each sample, more than fifteen *E. coli* clones were picked up for sequencing. The results were analyzed by DNA methylation analysis platform (http://services.ibc.uni-stuttgart.de/BDPC/BISMA/).

### RNA extraction and quantitative real-time PCR

The total RNA of high quality was isolated from cells and tissues of embryos (E8.5, E9.5, E12.5 and E18.5) using TRIzol™ Reagent (Thermo Scientific™). To synthesize high-quality cDNAs for real-time PCR, a mount of 500 ng of total RNA was directly used as template with ReverTra Ace® qPCR RT Master Mix (TOYOBO). To obtain high-quality RNA from embryos with low-cell number (E3.5, E6.5 and E7.5), samples were lysed in 150 μl of 4 M guanidine isothiocyanate solution (Thermo Scientific™) at 42 ℃ for 10 min. Total RNA pellets were concentrated by UltraPure™ Glycogen (Thermo Scientific™). To get enough amount of template for real-time PCR, two rounds of amplification were performed according to a reported protocol (Chen et al., 2017). Quantitative real-time PCR was performed on Bio-Rad CFX96 using the SYBR^®^ qPCR Mix (TOYOBO) in triplicate. All gene expression was calculated based on the 2^−∆∆Ct^ method after normalization to the transcript level of the internal standard gene, *Gapdh*. All the primer sequences are listed in (Table S1).

### RNA sequencing and analyzing

To separate ICM and TE from E3.5 embryos(Liu et al., 2016), blastocysts were broken using micromanipulator and washed three times in HCZB. The zona pellucidae of blastocysts were digested with 0.5% pronase E (Sigma) for 10-15 min and washed three times again in HCZB. The embryos were then incubated in Ca^2+^-free CZB for 30 min in 37 ℃, and the ICM and TE were separated into single cells by gently pipetting using a pipette with a diameter of 40–60 μm. Each sample was obtained about 10 cells from one embryo for amplification according the cell shape. ICM cells are small and out-of-shape, while TE cells are big and smooth. The cDNAs of low-input cells were harvested according previously protocols (Picelli et al., 2014). The cDNA libraries were prepared by SPAPK DNA Sample Pre Kit (Enzymatics) for sequencing using an Illumina HiSeq X Ten sequenator. Total RNA was isolated from mouse placentas using Dynabeads™ mRNA DIRECT™ Purification Kit (Thermo Scientific™). RNA-seq libraries were prepared by VAHTS mRNA-seq V3 Library Prep Kit for Illumina® (Vazyme), and then applied for deep sequencing on Illumina NovaSeq6000 platform. All RNA-seq reads were mapped to mm9 with TopHat (version 2.2.1). The mapped reads were further analyzed by Cufflinks and the expression levels for each transcript were quantified as Fragments Per Kilobase of transcript per Million mapped reads (FPKM).

### Intracytoplasmic AG-haESCs and sperms injection

To generate SC embryos, AG-haESCs were treated with 0.05 μg/ml Demecolcine solution (Sigma) for 10-12 h and synchronized to M phase. These cells were trypsinized and suspended in HCZB medium. MII oocytes were collected from oviducts of superovulated B6D2F1 females (8-week-old). AG-haESCs were injected into the cytoplasm of MII oocytes in a droplet of HCZB medium containing 5 µg/ml cytochalasin B (Sigma) using a Piezo-drill micromanipulator. The reconstructed oocytes were cultured in CZB medium for 30 min and then activated for 5–6 h in Ca^2+^ free CZB with SrCl_2_. For ICSI experiments, the procedures were the same as described above, except that AG-haESCs were replaced by sperm. Following activation, all of the reconstructed embryos were cultured for 24 h in AA-KSOM (Merk) medium at 37 ℃ under 5% CO_2_ in air.

### Embryo transfer and cesarean section

Reconstructed embryos were cultured in AA-KSOM medium until the two-cell or blastocyst stage. Thereafter, 14-16 (for ICSI) and 18-20 (for ICAHCI) two-cell embryos were transferred into each oviduct of pseudopregnant ICR females at 0.5 days post copulation. The post-implantation embryonic development (from E6.5 to E18.5) was assessed by an autopsy, and the embryos were quickly removed from the uteri and photographed immediately. Some E12.5 and E18.5 embryos and placentas were weighed after being dried with absorbent paper.

### Placenta histology analysis

The placentas of E12.5 were fixed in 4 % PFA overnight at room temperature, dehydrated with gradient concentration of alcohol, and embedded in paraffin and sectioned. Serial sections (4 μm) prepared in the cross planes were mounted on slides and dewaxed in xylene and rehydrated in different concentration of alcohol, followed by staining with hematoxylin and eosin. In each placenta, five sections of different slides were picked up to calculate the total placenta area, decidual zone area, labyrinthine zone area, and basal zone area. The values were measured with ImageJ software, and the area is the average of five different sections of each sample.

### Collection of mature oocytes and measure of luteinizing hormone (LH)

Control (ICSI), DKO and TKO female mice (4-week old) were superovulated with 4 international units of pregnant mare’s serum gonadotropin (PMSG) for 48 h and then human chorionic gonadotropin (hCG) for 12 h. Mature oocytes were collected from ampulla of the fallopian tube after superovulation. Plasma was collected from 4-month female mice using heparin as an anticoagulant. All samples were centrifuged for 15 min at 1000 g at 4 °C within 30 min and the supernatant was collected for the next assay. LH signals were detected from plasma using Mouse LH / Luteinizing Hormone ELISA Kit (LSBio), following the manufacturer’s instructions.

### Statistical analysis

For all figures, results shown are the average mean ± standard error of the mean (s.e.m.). Statistical testing was performed using GraphPad Prism 7. Significance was determined by two-tailed, unpaired Student’s t-test. \**P* < 0.05, \*\**P* < 0.01, \*\*\**P* < 0.001, n.s means no significant difference.

### Data availability

The RNA sequencing data of blastocysts and placentas from this study have been deposited in the Gene Expression Omnibus (GEO) under accession code GSE132254. The RNA sequencing data of AGH-OG3 and *H19*^ΔDMR^*-IG*^ΔDMR^-AGH-OG3 obtained from our previous work under the accession number GSE60072. The RNA sequencing data of ICM and TE of E3.5 and Epi and VE of E6.5 from previous work under the accession number GSE76505. All other data supporting the findings of this study are available from the corresponding author on reasonable request.

## Compliance and ethics

The authors declare that they have no conflict of interest.

## Acknowledgements

This study was partly supported by Genome Tagging Project, Fountain-Valley Life Sciences Fund of University of Chinese Academy of Sciences Education Foundation and grants from the Chinese Academy of Sciences (XDB19010204, OYZDJ-SSW-SMC023 and Facility-based Open Research Program), the National Natural Science Foundation of China (31530048, 81672117, 31730062, 31821004 and 31601163), and Shanghai Municipal Commission for Science and Technology (16JC1420500, 17JC1420102, 17JC1400900 and 17411954900).

## Author contributions

J.L. conceived of the project. Q.L. and Q.Y. performed the ICAHCI experiments. Y.L. and K.W. analyzed RNA-seq data. Q.L. and S.H. performed cell culture, histomorphology and molecular biological experiments. L.Z., W.L. and B.C. helped with embryo transplantation and the mouse experiments. Q.L., Y.L. and J.L. designed the experiments, analyzed the data and wrote the paper. All authors contributed to the manuscript.

## SUPPLEMENTAL FIGURES

**Figure S1.**
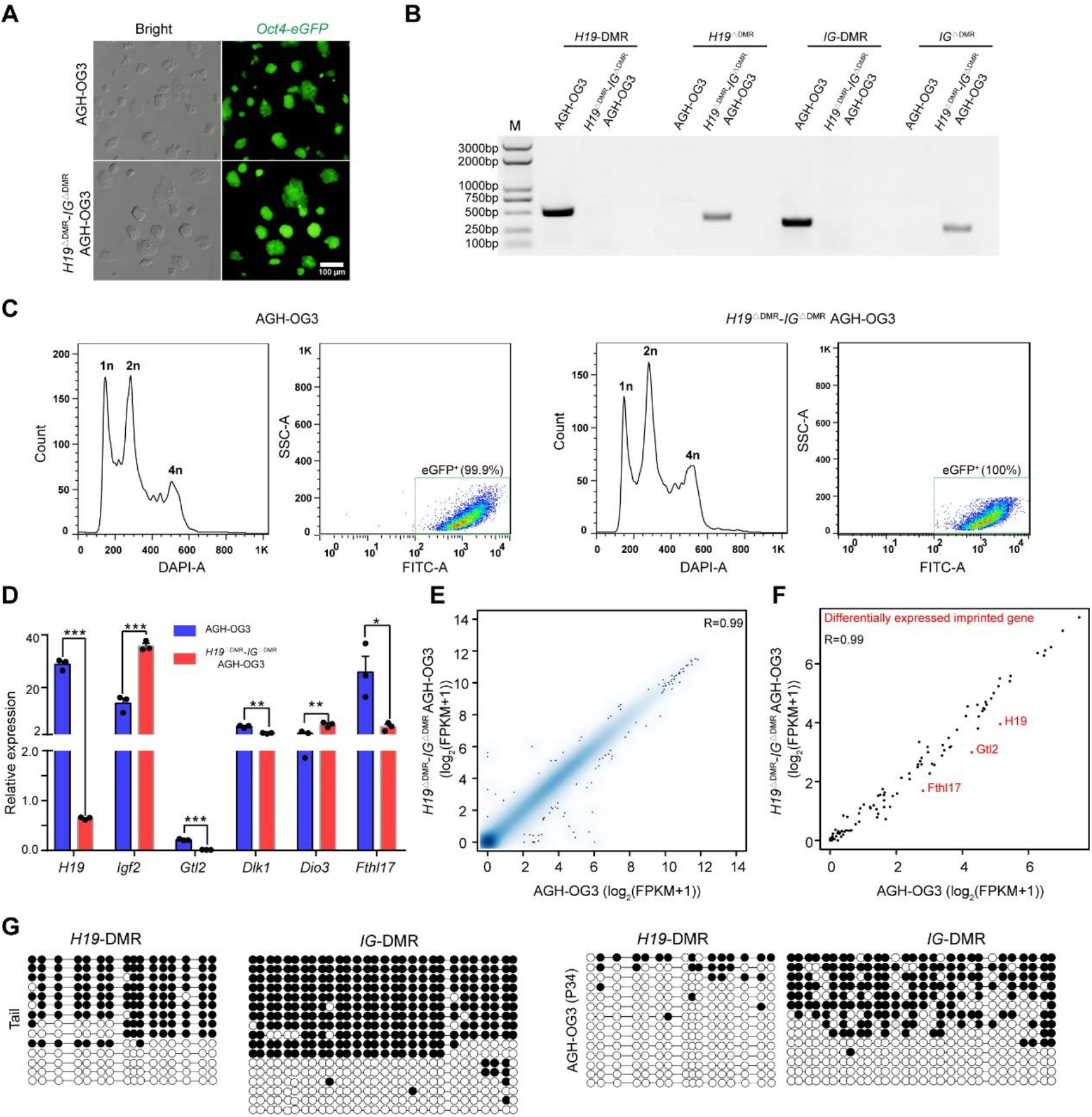
Characterizations of WT and DKO AG-haESCs. Related to Figure 1. (A) Images of WT AG-haESC (AGH-OG3) and DKO-AG-haESC (*H19*^ΔDMR^-*IG*^ΔDMR^-AGH-OG3) lines carrying an *Oct4-eGFP* transgene. Experiments were repeated three times independently, with similar results. (B) Genotyping analysis of *H19*^ΔDMR^-*IG*^ΔDMR^-AGH-OG3 (DKO-AG-haESC) lines, AGH-OG3 (WT-AG-haESCs) as the control sample. M, DNA marker. (C) FACS-mediated haploid cell-enrichment. (D) Transcriptional analysis of imprinted genes, *H19*, *Igf2*, *Gtl2*, *Dlk1*, *Dio3* and *Fthl17* in AGH-OG3 and *H19*^ΔDMR^-*IG*^ΔDMR^-AGH-OG3 cells. The expression values were normalized to that of *Gapdh*. Data are shown as the average mean ± s.e.m. \**P* < 0.05, \*\**P* < 0.01, \*\*\**P* < 0.001. (E) Genome-scale transcriptional similarity between WT and DKO AG-haESCs. The gene expression level was measured as log_2_(FPKM+1). The Pearson correlation coefficient (R) is 0.99. (F) The scatter plot of imprinted gene expression from WT-AG-haESC and DKO-AG-haESCs. The gene expression level was measured as log_2_(FPKM+1). The Pearson correlation coefficient (R) is 0.99. (G Methylation analysis of the *H19* and *IG*-DMRs in tail (C57BL/6 female) and AGH-OG3. Open circles represent unmethylated CpG sites, whereas filled circles represent methylated CpG sites.

**Figure S2.**
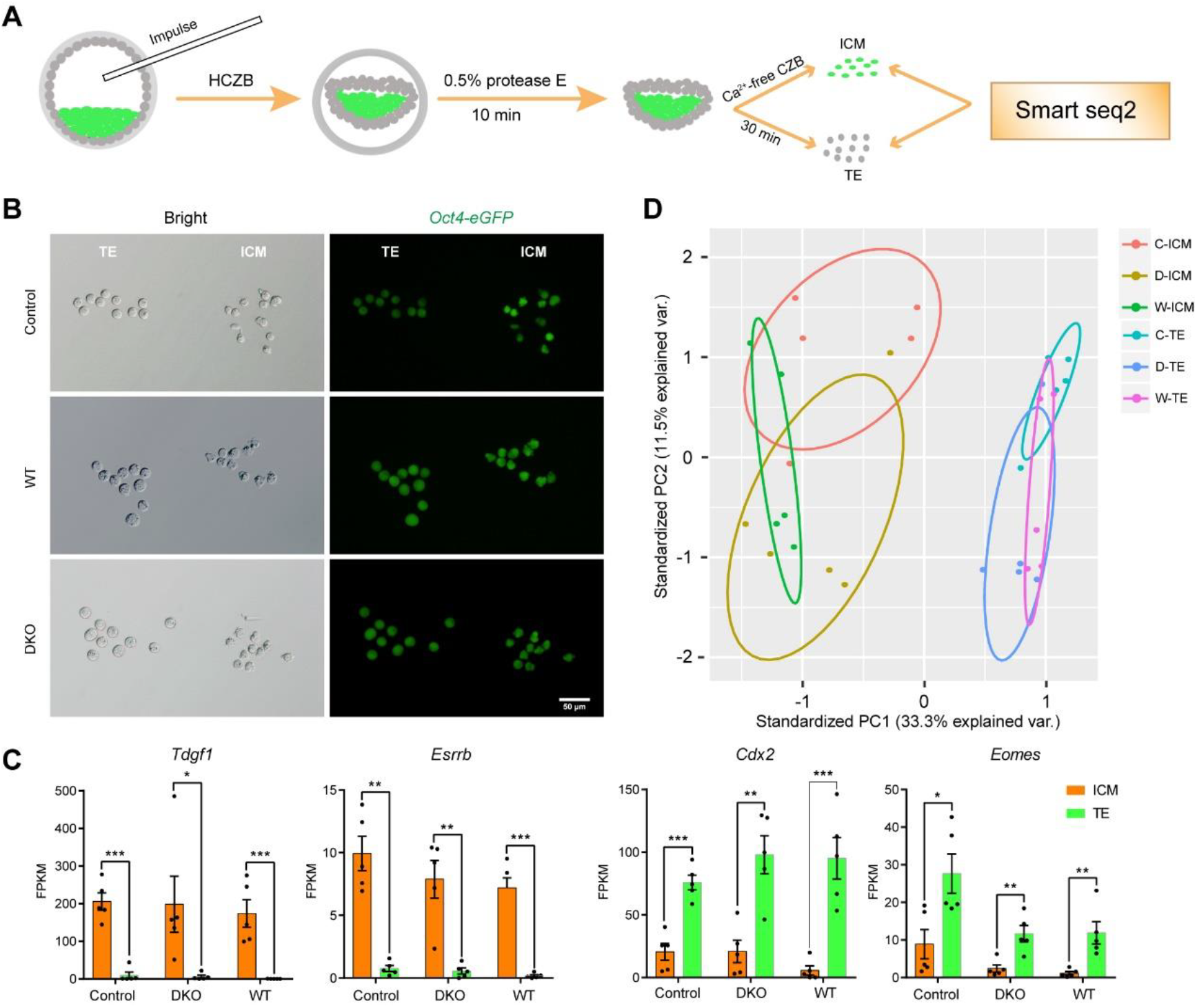
RNA-seq analysis of SC blastocysts. Related to Figure 1. (A) Workflow of isolation of inner cell mass (ICM) and trophectoderm cell (TE) and RNA sequencing through Smart-seq2. (B) Images showing that TE and ICM cells isolated from control (ICSI), WT and DKO SC blastocysts both carrying *eGFP* signals. TE cells are big and smooth; ICM cells are small and out-of-shape. (C) The bar plot showing the expression level of ICM marker genes (*Tdgf1* and *Esrrb*) and TE marker genes (*Cdx2* and *Eomes*) in the isolated ICM and TE based on RNA-seq data. Data are shown as the average mean ± s.e.m. \**P* < 0.05, \*\**P* < 0.01, \*\*\**P* < 0.001. (D) The principle component analysis (PCA) of RNA-seq data from ICM and TE obtained through oocyte injection of sperm (C), WT (W) and DKO (D) AG-ESCs respectively.

**Figure S3.**
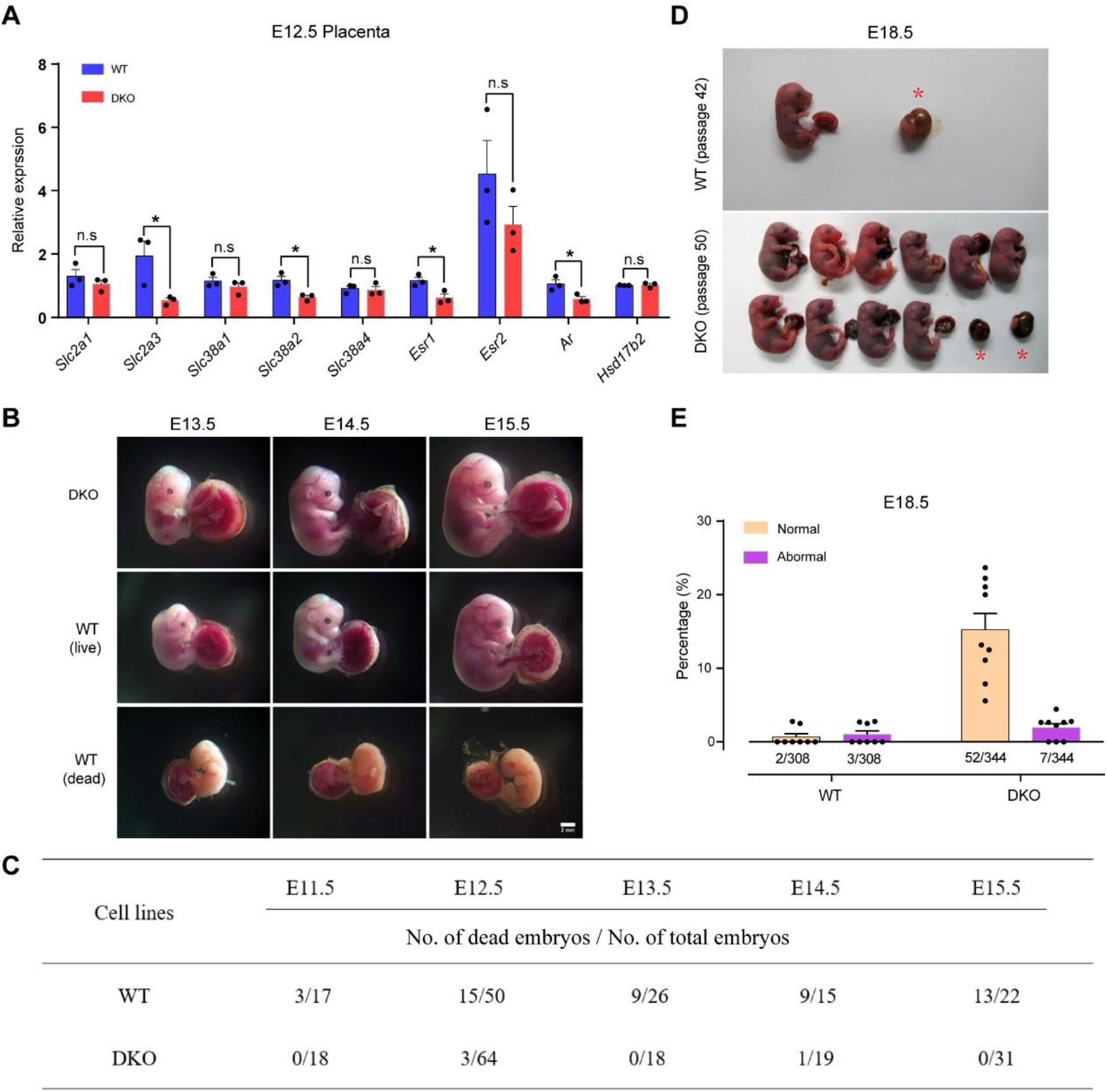
*H19*-DMR and *IG*-DMR deletions (DKO) enhance the SC embryonic development. Related to Figure 3. (A) Transcriptional analysis of glucose transporter genes (*Slc2a1* and *Slc2a3*), system A amino acid transporter genes (*Slc38a1*, *Slc38a2*, and *Slc38a4*), estrogen receptor (*Esr1* and *Esr2*), androgen receptor (*Ar*) and the inactivator of testosterone and estrogen (*Hsd17b2*) in E12.5 placentas of WT and DKO (*n* = 3 placentas for each group). (B) Representative images of SC fetuses derived from DKO (top) and WT (middle and bottom) haploid cells. Experiments were repeated five times independently, with similar results. (C) Summary of the number of dead embryos (E11.5 to E15.5) in all obtained SC embryos derived from WT and DKO AG-haESCs. Numerator means the dead embryos and denominator means the total number of embryos. (D) Images of SC embryos on E18.5. Red asterisks show degenerative embryos. (E) The ratios of normal and abnormal SC embryos (E18.5) derived from WT (*n* = 8 recipients) and DKO (*n* = 9 recipients) haploid cells. Numbers under the bars indicate E18.5 embryos/transferred two-cell embryos. Data are shown as the average mean ± s.e.m in A and E. \**P* < 0.05. n.s means no significant difference.

**Figure S4.**
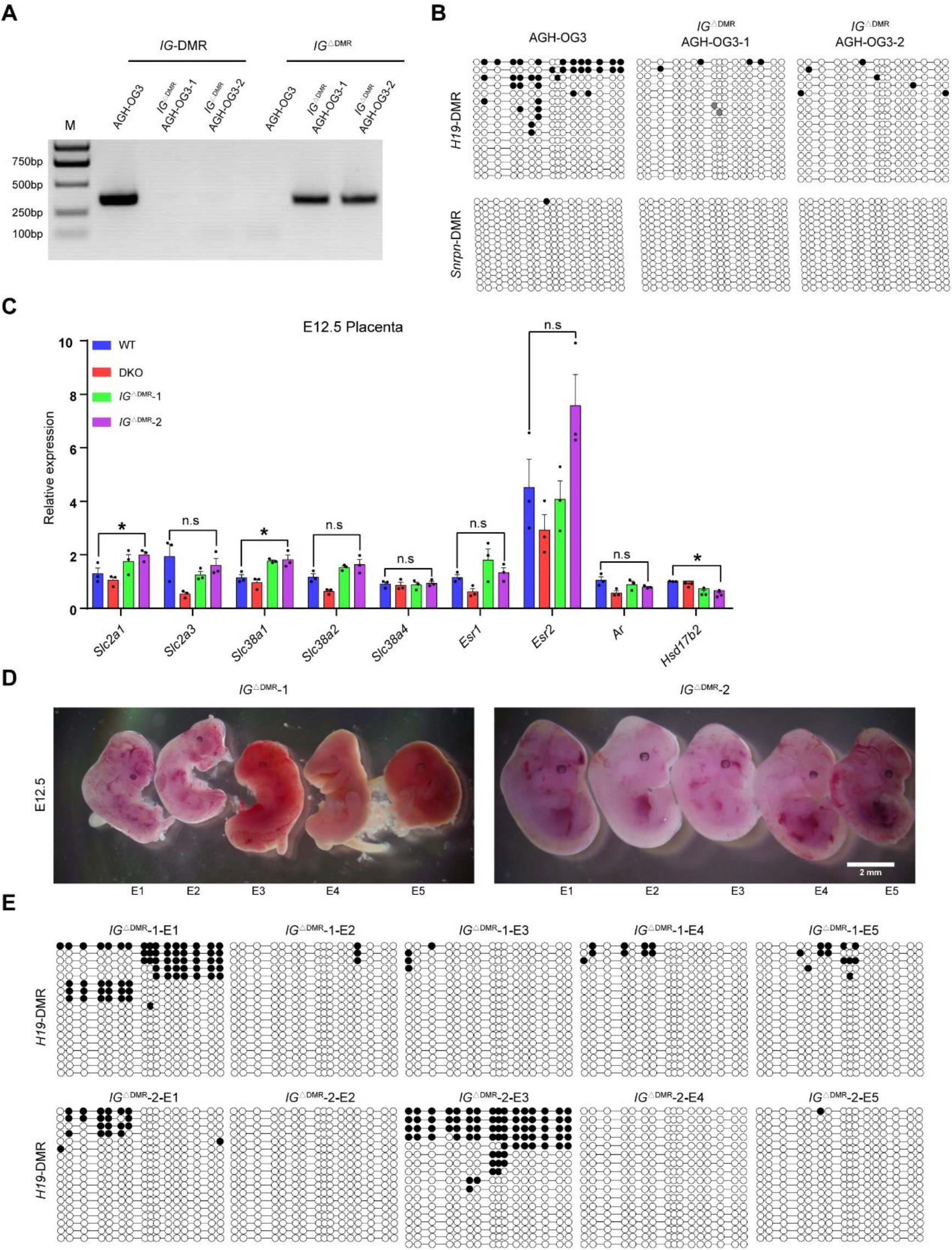
Abnormal development of SC embryos derived from *IG*^ΔDMR^-AG-haESCs. Related to Figure 4. (A) Genotyping analysis of two *IG*^ΔDMR^-AGH-OG3 cell lines. (B) Methylation analysis of the *H19* and *Snrpn-*DMRs in AGH-OG3 and *IG*^ΔDMR^-AGH-OG3. Open circles represent unmethylated CpG sites, whereas filled circles represent methylated CpG sites. (C) Transcriptional analysis of glucose transporter genes (*Slc2a1* and *Slc2a3*), system A amino acid transporter genes (*Slc38a1*, *Slc38a2*, and *Slc38a4*), estrogen receptor (*Esr1* and *Esr2*), androgen receptor (*Ar*) and the inactivator of testosterone and estrogen (*Hsd17b2*) in E12.5 placentas of WT, DKO and *IG*^ΔDMR^ (*n* = 3 placentas for each group). Data are shown as the average mean ± s.e.m. \**P* < 0.05. n.s means no significant difference. (D and E) Representative mages of *IG*^ΔDMR^ SC embryos on E12.5 (D) and methylation analysis of the *H19-*DMR in these embryos (E). Open circles represent unmethylated CpG sites, whereas filled circles represent methylated CpG sites.

**Figure S5.**
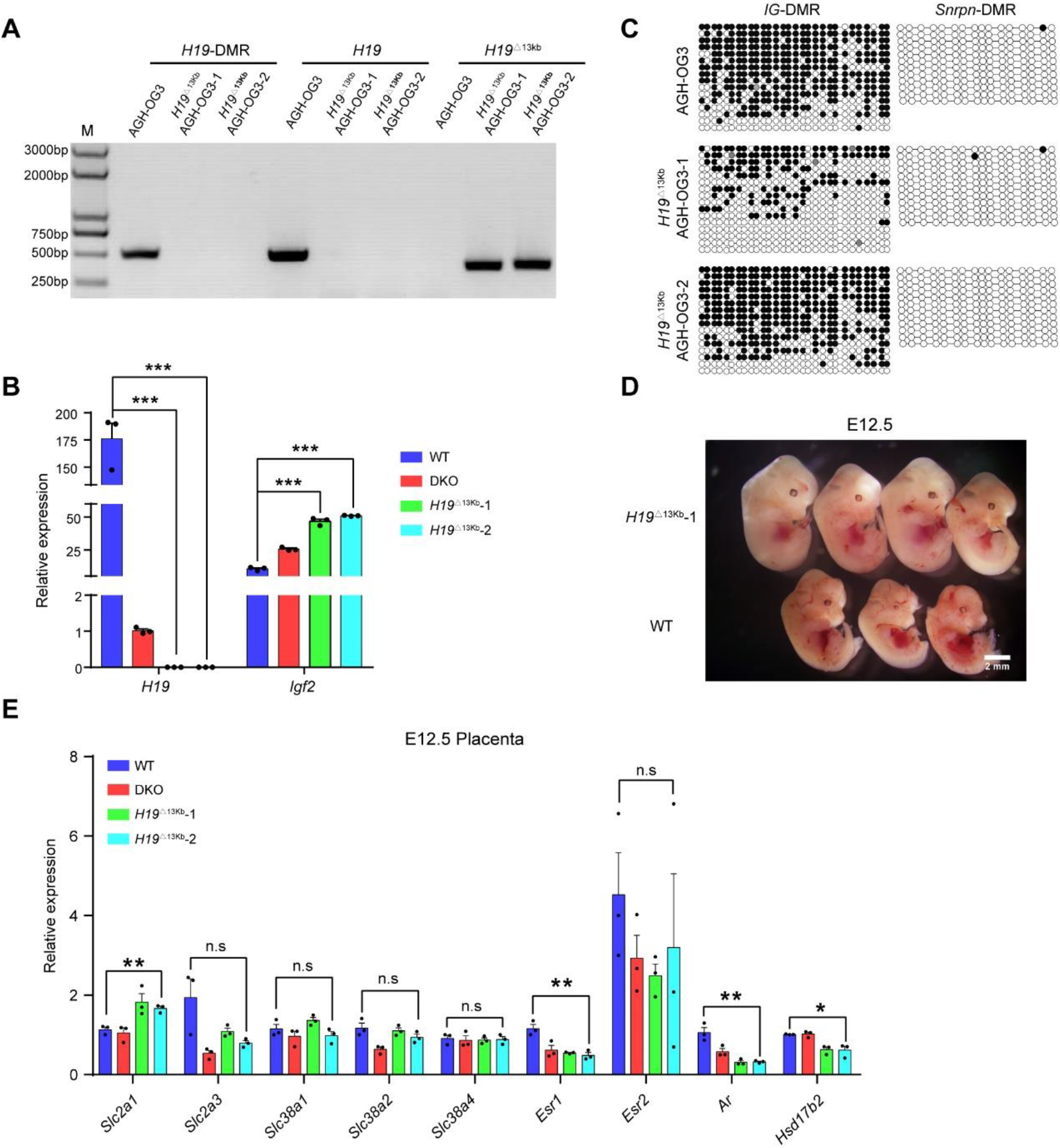
Characterizations of *H19*^Δ13kb^-AG-haESCs. Related to Figure 5. (A) Genotyping analysis of two *H19*^Δ13kb^-AGH-OG3 cell lines, AGH-OG3 as the control sample. M, DNA marker. (B) Transcriptional analysis of *H19* and *Igf2* genes in AGH-OG3, *H19*^ΔDMR^-*IG*^ΔDMR^-AGH-OG3 and *H19*^Δ13kb^-AGH-OG3 cells, showing no *H19* expression in *H19*^Δ13kb^ AG-haESCs. The expression values were normalized to that of *Gapdh*. (C) Methylation analysis of the *IG* and *Snrpn*-DMRs in *H19*^Δ13kb^-AGH-OG3 and AGH-OG3. Open circles represent unmethylated CpG sites, whereas filled circles represent methylated CpG sites. (D) Images of *H19*^Δ13kb^ and WT SC embryos on E12.5. Experiments were repeated three times independently, with similar results. (E) Transcriptional analysis of glucose transporter genes (*Slc2a1* and *Slc2a3*), system A amino acid transporter genes (*Slc38a1*, *Slc38a2*, and *Slc38a4*), estrogen receptor (*Esr1* and *Esr2*), androgen receptor (*Ar*) and the inactivator of testosterone and estrogen (*Hsd17b2*) in E12.5 placentas of WT, DKO and *H19*^Δ13kb^ (*n* = 3 placentas for each group). Data are shown as the average mean ± s.e.m in B, E. \**P* < 0.05, \*\**P* < 0.01, \*\*\**P* < 0.001. n.s means no significant difference.

**Figure S6.**
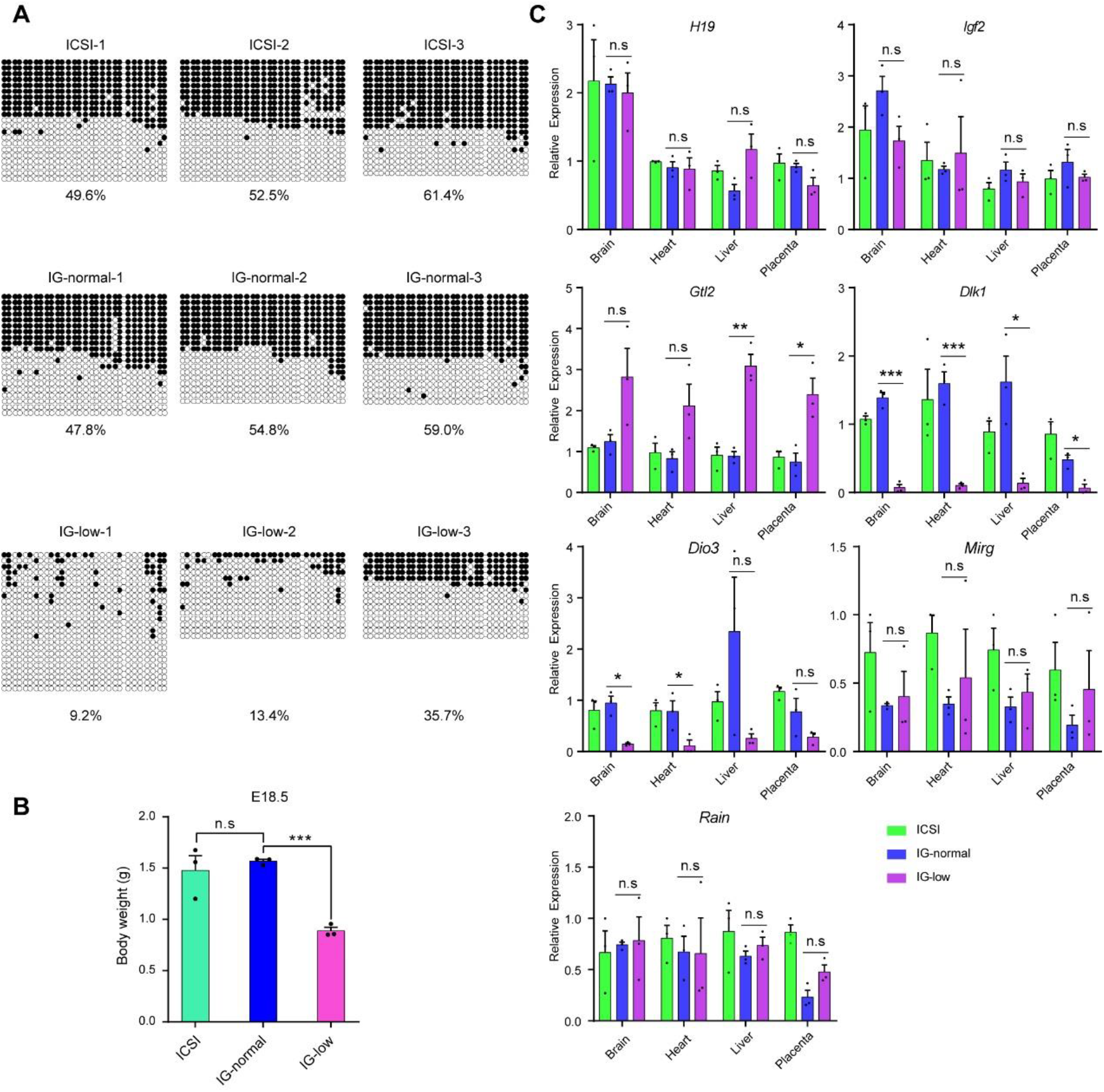
The correlation between hypomethylation of *IG*-DMR and growth retardation of SC embryos from *H19*^Δ13kb^-AGH-OG3 cells at late-gestation. Related to Figure 5. (A) Methylation analysis of *IG*-DMR on E18.5 embryos derived from sperm (ICSI) and *H19*^Δ13kb^-AGH-OG3. *n* = 3 embryos for each group. IG-normal SC embryos carrying normal methylation and IG-low SC embryos with hypomethylation compare with ICSI. Open circles represent unmethylated CpG sites, whereas filled circles represent methylated CpG sites. (B) Body weight of E18.5 embryos derived from sperm (ICSI) and *H19*^Δ13kb^-AGH-OG3 (IG-normal and IG-low), showing that IG-low embryos exhibit growth retardation. (C) Transcriptional analysis of imprinted genes *H19*, *Igf2*, *Gtl2*, *Dlk1*, *Dio3*, *Mirg* and *Rain* in different tissues of E18.5 pups (ICSI, IG-normal and IG-low), including brain, heart, liver and placenta (*n* = 3 samples for each group). The expression values were normalized to that of *Gapdh*. Data are shown as the average mean ± s.e.m in B and C. \**P* < 0.05, \*\**P* < 0.01, \*\*\**P* < 0.001. n.s means no significant difference.

**Figure S7.**
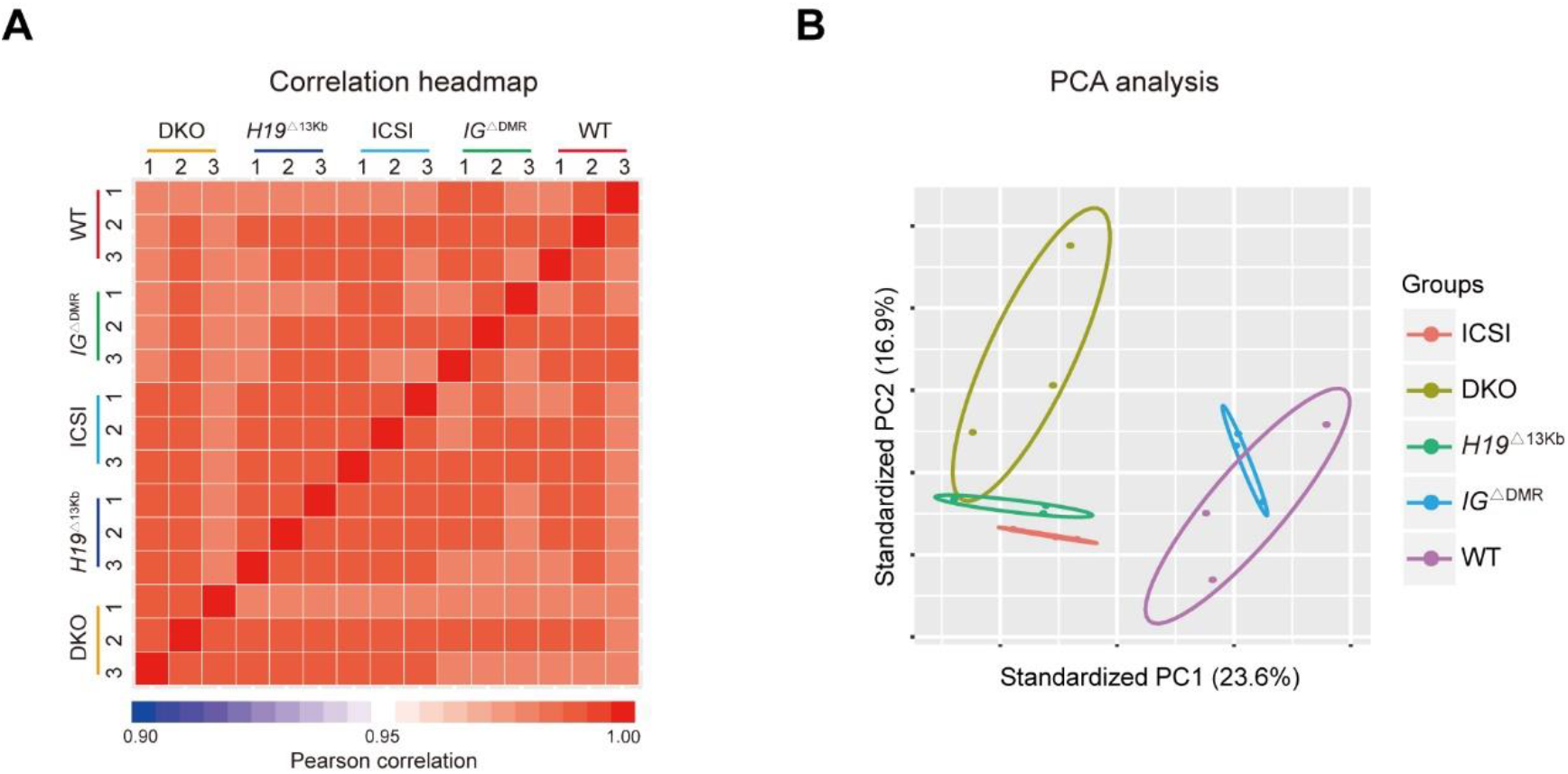
RNA-seq analysis of placentas with different deletions. Related to Figure 6. (A) The correlation heatmap of RNA dataset from placentas obtained through oocyte injection of sperm, WT, DKO, *IG*^ΔDMR^, and *H19*^Δ13kb^ AG-haESCs respectively. (B) The principle component analysis (PCA) based on all expressed genes of RNA-seq data.

**Figure S8.**
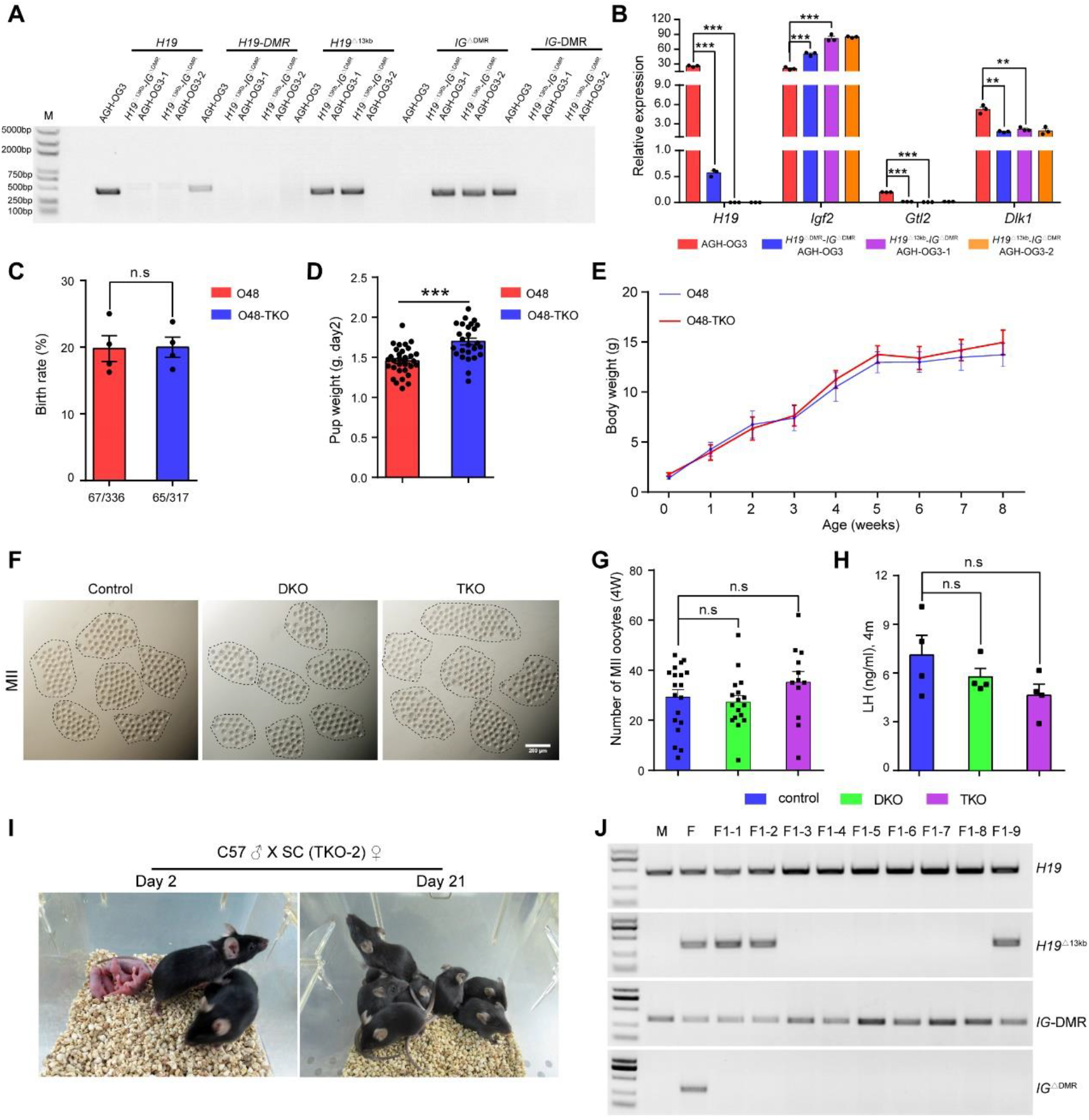
Characterizations of TKO AG-haESCs. Related to Figure 7. (A) Genotyping analysis of two *H19*^Δ13kb^-*IG*^ΔDMR^-AGH-OG3 cell lines (TKO-AG-haESCs), AGH-OG3 as the control sample. M, DNA marker. (B) Transcriptional analysis of *H19*, *Igf*2, *Gtl2* and *Dlk1* genes in AGH-OG3, *H19*^ΔDMR^-*IG*^ΔDMR^-AGH-OG3 and *H19*^Δ13kb^-*IG*^ΔDMR^-AGH-OG3 (two different cell lines). The expression values were normalized to that of *Gapdh*. (C) The birth rate of newborn SC pups from ICAHCI of O48 and O48-TKO cells. The O48 cell line was described in previous work (termed also as *H19*^ΔDMR^-*IG*^ΔDMR^-AGH-2) (Zhong et al., 2015). (D) Body weight of O48-TKO SC pups are significantly higher than that of O48 on day 2 after birth. (E) Growth curve of O48 (*n* = 10) and O48-TKO SC mice (*n* = 11), showing comparable growth rate between DKO and TKO SC pups. (F) Images of mature oocytes obtained from superovulated 4-week old females, including ICSI (control), DKO and TKO mice. The dotted lines indicate oocytes from one female. (G) Quantification of mature oocytes derived from ICSI (*n* = 19 females), DKO (*n* = 17 females) and TKO (*n* = 12 females) groups. (H) Measurement of luteinizing hormone (LH) in plasma from 4-month mice (ICSI, DKO and TKO). *n* = 4 mice for each group. Data are shown as the average mean ± s.e.m in B-D, G and H. \*\**P* < 0.01, \*\*\**P* < 0.001. n.s means no significant difference. (I) Adult TKO SC mice and their offspring. Experiments were repeated three times independently, with similar results. (J) Genotyping analysis of the *H19*, *H19*-DMR and *IG*-DMR deletions in the alive progeny of SC TKO mice.

**Table S1.**
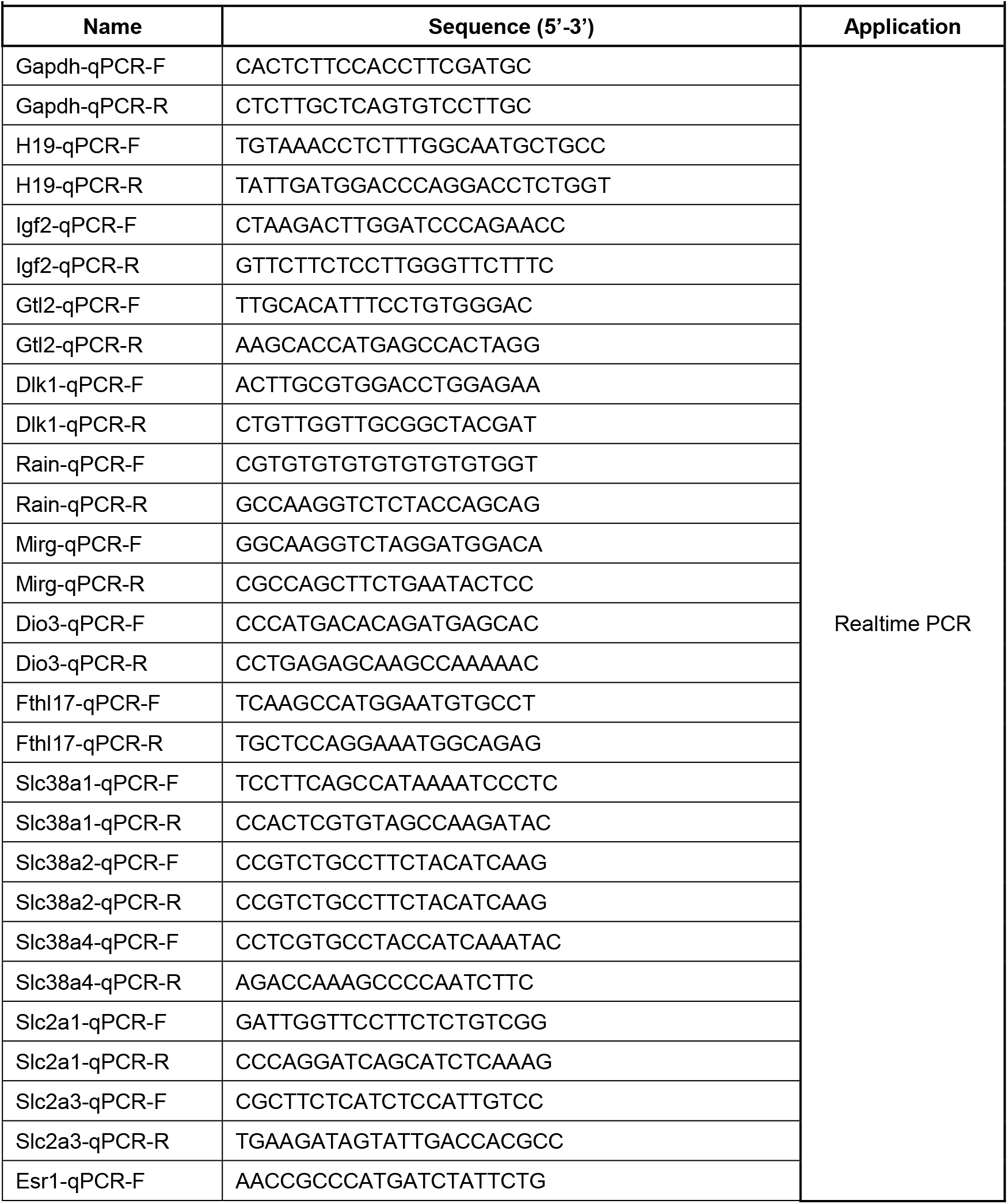
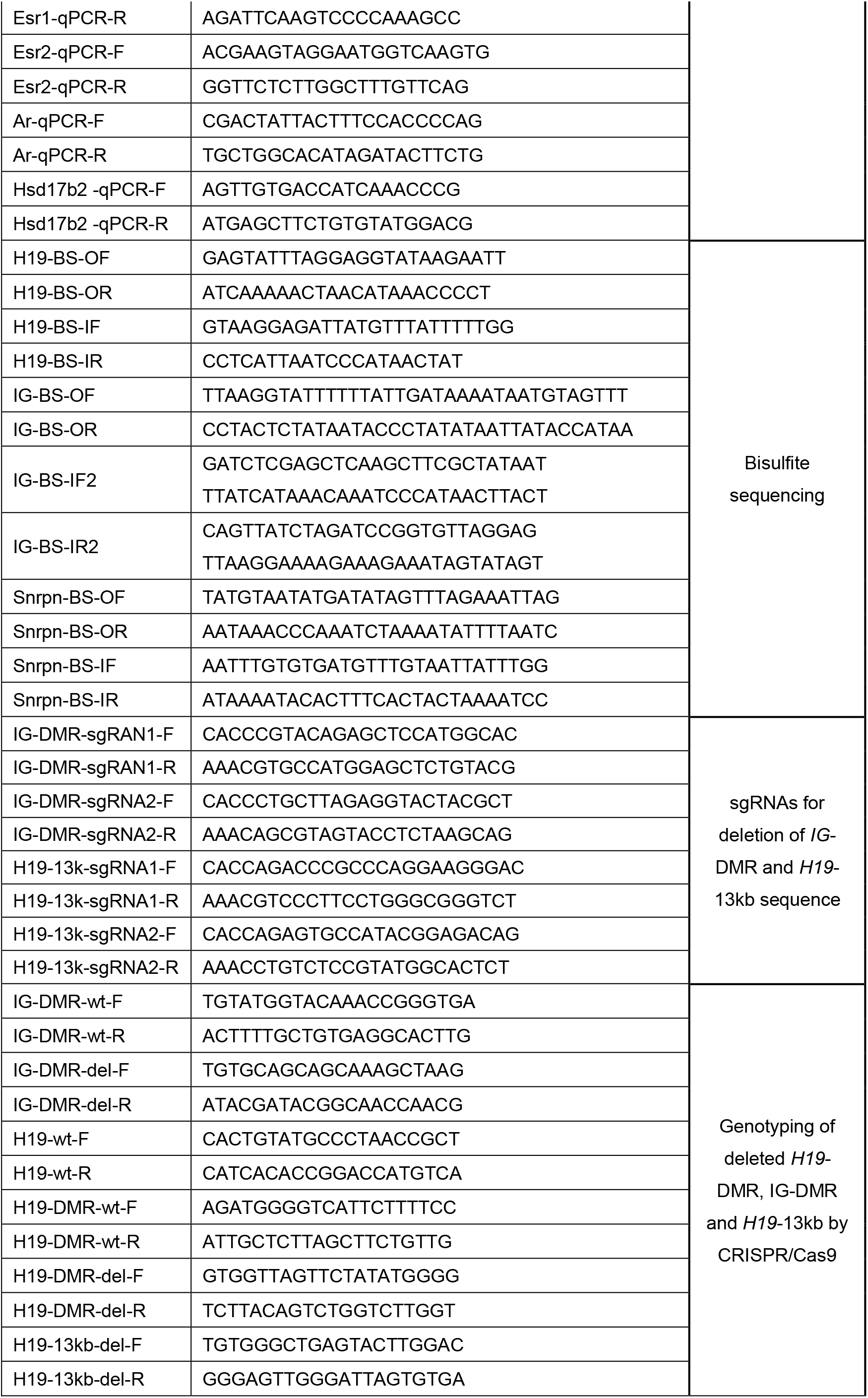
List of primer and sgRNA sequence information, related to experimental procedures

